# Structural Insights into the Unexpected Agonism of Tetracyclic Antidepressants at Serotonin Receptors 5-HT_1e_R and 5-HT_1F_R

**DOI:** 10.1101/2023.10.05.561100

**Authors:** Gregory Zilberg, Alexandra K. Parpounas, Audrey L. Warren, Bianca Fiorillo, Davide Provasi, Marta Filizola, Daniel Wacker

## Abstract

Serotonin (5-hydroxytryptamine, 5-HT) acts via 13 different receptors in humans. Of these receptor subtypes, all but 5-HT_1e_R have confirmed roles in native tissue and are validated drug targets. Despite 5-HT_1e_R’s therapeutic potential and plausible druggability, the mechanisms of its activation remain elusive. To illuminate 5-HT_1e_R’s pharmacology in relation to the highly homologous 5-HT_1F_R, we screened a library of aminergic receptor ligands at both receptors and observe 5-HT_1e/1F_R agonism by multicyclic drugs described as pan-antagonists at 5-HT receptors. Potent agonism by tetracyclic antidepressants mianserin, setiptiline, and mirtazapine suggests a mechanism for their clinically observed anti-migraine properties. Using cryoEM and mutagenesis studies, we uncover and characterize unique agonist-like binding poses of mianserin and setiptiline at 5-HT_1e_R distinct from similar drug scaffolds in inactive-state 5-HTR structures. Together with computational studies, our data suggest that these binding poses alongside receptor-specific allosteric coupling in 5-HT_1e_R and 5-HT_1F_R contribute to the agonist activity of these antidepressants.

## Introduction

Serotonin (5-Hydroxytryptamine, 5-HT) is a conserved neurotransmitter that coordinates the physiological parameters of diverse processes across organ systems ranging from memory, mood and sleep to gastrointestinal motility and vasodilation (*1, 2*). In humans, the effects of serotonin are mediated by 13 5-HT receptors (5-HTRs), 12 of which are G protein-coupled receptors (GPCR) whereas 5-HT_3_Rs comprise a family of pentameric ligand-gated ion channels (*1*). Due to serotonin’s important role in human health and disease inside and outside the CNS, 5-HTRs are targeted by numerous medications and illicit drugs, and thus represent major drug targets for numerous conditions (*3*). For instance, amongst the 5-HT_1_R subfamily, 5-HT_1A_R is targeted by antidepressants and anxiolytics such as buspirone and vilazodone, while 5-HT_1B_R, 5-HT_1D_R, and to a lesser extent 5-HT_1F_R, are targeted by anti-migraine triptan medications such as sumatriptan and frovatriptan (*4, 5*). In striking contrast, among 5-HTRs, 5-HT_1e_R stands alone in its designation as an orphan receptor due to no reports, to date, characterizing its physiological role (indicated by the International Union of Basic and Clinical Pharmacology with a lowercase appellation) (*1*). This is the result of a lack of 5-HT_1e_R-selective compounds, studies on the receptor’s pharmacology and molecular mechanisms, and, critically, a genetically accessible rodent orthologue for knockout experiments (*6–8*). For instance, the nominally 5-HT_1e_R-selective high affinity agonist BRL-54443 (*6*) exhibits equal or superior potency at the highly homologous receptor 5-HT_1F_R (*7*). Like 5-HT_1e_R, 5-HT_1F_R is also an understudied 5-HTR. However, 5-HT_1F_R has been implicated in migraine pathophysiology, and lasmiditan (Reyvow), a 5-HT_1F_R-selective drug, has recently been approved for treating migraines (*9*). In fact, 5-HT_1e_R and 5-HT_1F_R are routinely left out of off-target testing in older publications; 5-HT_1F_R in particular is not routinely tested by the NIH psychoactive drug screening program in their standard binding and cAMP inhibition panels (*10*). As a result, potential activities of known serotonergic drugs at these enigmatic receptors remain largely unknown, even though many of these are “pan-aminergic” drugs that are known to possess high affinity at multiple 5-HTR subtypes. While this oversight contributes to our poor understanding of 5-HT_1e_R’s and 5-HT_1F_R’s pharmacological and molecular mechanisms, examining the activity of serotonergic drugs may uncover physiological (side) effects of drugs mediated by these receptors. Moreover, such activity could potentially be harnessed to explore pharmacophores that yield receptor-selective probes, such as first in class 5-HT_1e_R-selective tool compounds with which to explore previous suggestions of 5-HT_1e_R as a viable pharmacological target in the treatment of certain types of cancers (*11, 12*).

To address these gaps in our molecular understanding of 5-HT_1e_R and 5-HT_1F_R pharmacology and function, we herein combine pharmacological assays, cryoEM structure determination, structure-activity relationship (SAR) studies, as well as computer simulations. Specifically, screening a focused library of aminergic compounds and drugs against both receptors revealed potent agonist activity by multiple compounds, including the multicyclic prescription drugs mianserin and pimethixene. We subsequently determined structures of 5-HT_1e_R signaling complexes activated by the antidepressants mianserin and setiptiline and performed molecular dynamics (MD) simulations of these structures embedded in an explicit lipid-water environment, as well as *in vitro* pharmacological assays. Our data reveal key insights into the binding of non-tryptamine agonists at 5-HT_1e_R and 5-HT_1F_R, provide insights into clinically reported side effects of mianserin and its analogs, and offer hints at how these supposedly pan-antagonistic compounds activate these enigmatic receptors.

## Results

### Curated library screening uncovers activity of aminergic medications and research chemicals at serotonin receptors 5-HT_1e_R and 5-HT_1F_R

Functional efficacies of drugs known to interact non-selectively with monoaminergic receptors are poorly characterized at 5-HT_1e_R and 5-HT_1F_R relative to other serotonin receptors. A closer examination of these interactions could provide additional insight into their physiological roles and potentially explain drug side effects.

We thus set out to characterize the effects of selected medications and research chemicals for aminergic GPCRs at both receptors to define their molecular properties and potentially obtain novel pharmacological leads for the development of future receptor-selective probes (Fig. 1, Fig. S1). Like the other members of the 5-HT_1_R family, 5-HT_1e_R and 5-HT_1F_R activation stimulates inhibitory G proteins such as G_i1-3_, G_o_, and G_z_, which inhibit adenylyl cyclase, thereby lowering cellular cAMP levels. To identify potential agonist activity of aminergic drugs at 5-HT_1e_R and 5-HT_1F_R, we measured receptor-mediated reduction in cellular cAMP using the GloSensor biosensor (*13*) (Fig. S1A). We also measured receptor-mediated arrestin recruitment (*1*) via the PRESTO-TANGO assay (*14, 15*), which served as an orthogonal assay to determine receptor-mediated activity in a distinct pathway (Fig. S1B). We first validated both assays using serotonin (*7, 16*), before assaying a panel of 87 select aminergic receptor ligands, including endogenous neurotransmitters such as serotonin and histamine, research chemicals, and clinically used drugs at 10 µM concentration (Fig. 1A, Table S1). We herein found a high number of agonist activities, as defined by an at least 4-fold (log2-fold change of 2) increase of signal over DMSO-treated controls: 64 compounds showed agonist activity in at least one pathway of either 5-HT_1e_R or 5-HT_1F_R (Fig. 1A-B, Table S1).

**Fig. 1.**
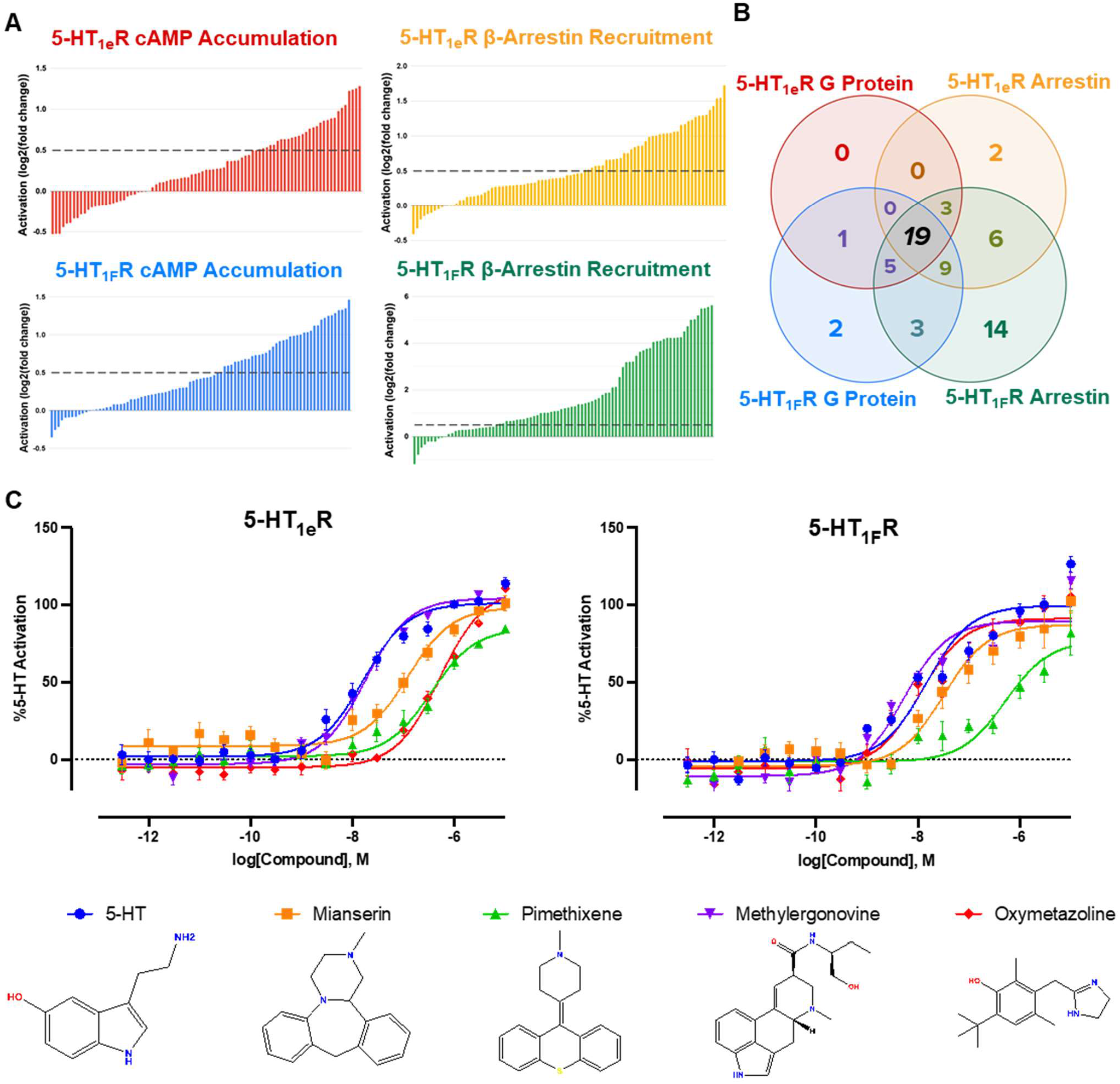
A curated screen of aminergic receptor ligands reveals 5-HT_1e_R and 5-HT_1F_R agonism by chemically diverse drugs and research chemicals. **A**, Activation of 5-HT_1e_R and 5-HT_1F_R as measured by cAMP accumulation and β-arrestin recruitment performed in HEK293T cells. Experiments were performed in quadruplicate using 10 µM ligand concentration and data is shown as log2-fold change with compounds considered active when above 0.5 (dashed line). β-arrestin recruitment data was normalized against DMSO, and cAMP accumulation data was normalized against effects in cells not transfected with receptor. **B**, Venn diagram showing the number of compounds active in a single or multiple screens. **C**, 2D chemical structures and respective concentration response curves in 5-HT_1e_R- or 5-HT_1F_R-mediated Gi1 activation in HEK293T cells. Experiments were performed in triplicate and mean±SEM from at least 2 (n=2) independent experiments were averaged and normalized to 5-HT’s response.

Of the compounds screened, 36 ligands activated both receptors across at least 3 screens (Fig. 1B, Table S1), including the canonical agonist 5-HT, the 5-HT_1e_R/5-HT_1F_R-selective agonist BRL-54443, and 2-Br-LSD in agreement with a recent report (*17*). 21 of these 36 compounds showed sub-10 µM affinity at either 5-HT_1e_R, 5-HT_1F_R, or both, in prior studies (*10, 17*), while no activity at the tested receptors has been reported for the remaining 15 compounds (Table S1; see “Screening Analysis” in methods for how active compounds were determined). Additionally, we observed that 3 compounds, the 5-HT_1A_R partial agonists buspirone and ipsapirone, and the antipsychotic raclopride showed significant activity in both 5-HT_1F_R screens without significant activity in the corresponding 5-HT_1e_R assays (Fig. 1B, Table S1).

To validate our results from the large scale screens and determine potencies and efficacies of a subset of these compounds, we performed concentration-response experiments using an orthogonal bioluminescent resonance energy transfer (BRET) G protein activation assay (*18*) (Fig. S1C). Of the compounds tested, we report nanomolar potencies for methylergonovine (5-HT_1e_R EC_50_=17.0 nM, 5-HT_1F_R EC_50_= 5.2 nM), oxymetazoline (5-HT_1e_R EC_50_=516.4 nM, 5-HT_1F_R EC_50_= 11.3 nM), pimethixene (5-HT_1e_R EC_50_=353.1 nM, 5-HT_1F_R EC_50_= 456.0 nM), and mianserin (5-HT_1e_R EC_50_=123.3 nM, 5-HT_1F_R EC_50_= 47.5 nM), while all other tested compounds exhibited weak-to-moderate µM potency agonism at both receptors (Fig. 1C, see Table S2 for standard error).

Studies previously reported 5-HT_1e_R and 5-HT_1F_R agonism of methylergonovine (Methergine, a second-line uterotonic and potential anti-migraine agent), while agonism of oxymetazoline (Afrin, a topical decongestant and alpha adrenergic receptor agonist) was previously only described for 5-HT_1F_R, 5-HT_1A_R, 5-HT_1B_R, and 5-HT_1D_R, but not 5-HT_1e_R (*19, 20*). By contrast, the potent agonism of pimethixene (Muricalm, an antihistamine and anticholinergic) and mianserin (Tolvon, a noradrenergic and specific serotonin antidepressant) were unexpected due to their reported antagonism for several GPCRs. Specifically, pimethixene is an antagonist at histamine, serotonin, and muscarinic acetylcholine receptors (*21, 22*), and mianserin is an antagonist at α-adrenergic and 5-HT_2_ receptors (*23, 24*), though weak partial agonism (EC_50_ = 530 nM) has been reported at the kappa opioid receptor (*25*).

### Functional characterization of the unexpected 5-HT_1e_R and 5-HT_1F_R agonism of tetracyclic antidepressants and structurally related drugs

Since mianserin’s high potency and agonism was particularly unexpected, we sought confirmation of its atypical activity at 5-HT_1e_R and 5-HT_1F_R. Indeed, several studies describe mianserin as a pan-aminergic, pan-serotonergic, or non-selective 5-HT_2_R and adrenergic α_2_R antagonist (*26–28*). We therefore decided to investigate the possible extension of this agonism to other similar human serotonin receptors using the previously described arrestin-recruitment assay (Fig. S2). In this assay, mianserin potently activated 5-HT_1e_R (EC_50_= 67 nM) with full efficacy, while it produced markedly weaker partial agonism at 5-HT_1F_R (EC_50_= 667 nM, Emax = 24%) and 5-HT_1D_R (EC_50_= 371 nM, Emax = 25%). In contrast, no noticeable agonism was observed at any of the other 5-HT receptors, with potential inverse agonism at 5-HT_2A_R and 5-HT_2B_R at high concentrations. Likewise, pimethixene-mediated arrestin recruitment at these receptors showed full agonism at 5-HT_1e_R (EC_50_= 453 nM), but no noticeable activity at the other receptors tested apart from high concentrations at 5-HT_1A_R (see Fig. S2 legend for standard errors). Together, we find that mianserin and pimethixene not only display selective agonism for 5-HT_1e_R and 5-HT_1F_R, but they also display distinct pharmacological activities at the two receptors. Specifically, both drugs appear to be balanced agonists in different pathways downstream of 5-HT_1e_R but they are biased towards G protein pathways at 5-HT_1F_R.

Given the atypical receptor agonism of the multicyclic drugs mianserin and pimethixene, scaffolds that typically antagonize aminergic GPCRs (*29*), we next systematically investigated the 5-HT_1e_R and 5-HT_1F_R activity of other compounds and clinically used medications with related scaffolds (Fig. 2). Among the tested compounds, several drugs including the muscle relaxant cyclobenzaprine (Amrix), as well as the antipsychotics chlorpromazine (Thorazine) and clozapine (Clozaril), display nanomolar potency at either 5-HT_1e_R or 5-HT_1F_R, albeit with distinct selectivities and efficacies. It should be noted that binding affinities, but not functional efficacies, of clozapine and chlorpromazine have been previously reported for 5-HT_1e_R and 5-HT_1F_R (*30*).

**Fig. 2.**
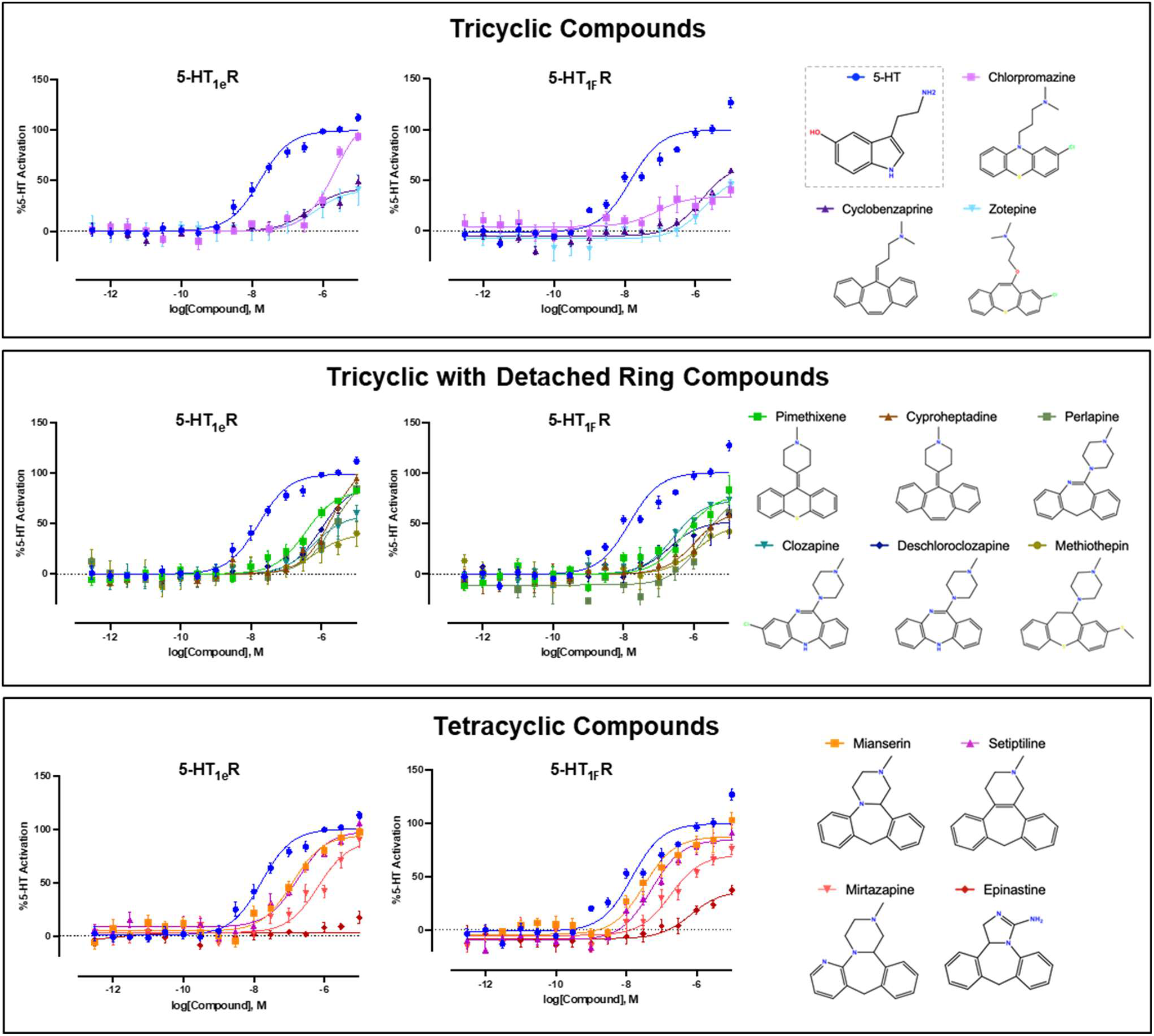
Activities of multicyclic drugs at 5-HT_1e_R and 5-HT_1F_R. 2D chemical structures and respective concentration response curves in 5-HT_1e_R- or 5-HT_1F_R-mediated cAMP accumulation in HEK293T cells are shown. Experiments were performed in triplicate and mean±SEM from at least two (n=2) independent experiments were averaged and normalized to 5-HT’s response.

These findings show that even multicyclic scaffolds distinct from that of mianserin can activate 5-HT_1e_R and 5-HT_1F_R. Using the aforementioned G protein BRET assay, we subsequently determined that setiptiline and mirtazapine, two tetracyclic antidepressants (TeCA) and closely related structural analogs of mianserin, exhibit the most potent and efficacious activity among the multicyclic drugs tested. Specifically, we find that setiptiline is a potent full agonist at both 5-HT_1e_R (EC_50_ = 171.0 nM) and 5-HT_1F_R (EC_50_ = 64.6 nM), whereas mirtazapine shows reduced potency at 5-HT_1e_R (EC_50_ = 1.04 μM) relative to 5-HT_1F_R (EC_50_ = 235.5 nM) (Fig. 2).

Together, these findings uncover unique aspects of 5-HT_1e_R’s and 5-HT_1F_R’s molecular pharmacology and illuminate previously underexplored facets of the polypharmacology of mianserin and related prescription drugs. Strikingly, mianserin and mirtazapine exhibit inadvertent anti-migraine effects in patients (*31, 32*), which could potentially be due to their herein found unexpected promiscuous agonism at the *bona fide* anti-migraine target 5-HT_1F_R.

### CryoEM studies of 5-HT_1E_R-Gα_i1_-Gβ_1_-Gγ_2_ complexes illuminate structural features of 5-HT_1e_R bound by tetracyclic antidepressants

Despite the clinical use of mianserin and related TeCAs, little is known about how they bind their cognate receptors, let alone how they activate 5-HT_1e_R and 5-HT_1F_R. To elucidate the molecular basis of these mechanisms, we next determined cryoEM structures of mianserin- and setiptiline-bound 5-HT_1e_R-G protein signaling complexes. This was done largely following previously published methodology (See Materials and Methods) (Fig. 3A, Fig. S3, S4, Table S3) (*33*). Briefly, 5-HT_1e_R containing a L111^3.41^W mutation (Superscripts denote Ballesteros– Weinstein Numbering (*34*)) and Gα_i1_β_1_γ_2_ (G_i1_) were separately expressed (*35*), subsequently combined in the presence of agonist, and complexes were isolated by gel-filtration chromatography. Peak fractions were concentrated and frozen on grids, and subsequently imaged and processed to obtain high-resolution cryoEM reconstructions. Mianserin- and setiptiline-bound 5-HT_1e_R-G_i1_ complex structures were determined at global resolutions of 3.30 Å and 3.32 Å, respectively, with local resolutions of ∼3.0 Å and ∼3.2 Å for the ligand binding pockets (Fig. S3, S4, Table S3). At these resolutions, we were able to unambiguously identify residue conformers within the ligand-binding pocket and elucidate mianserin and setiptiline’s binding poses and drug-receptor interactions.

**Fig. 3.**
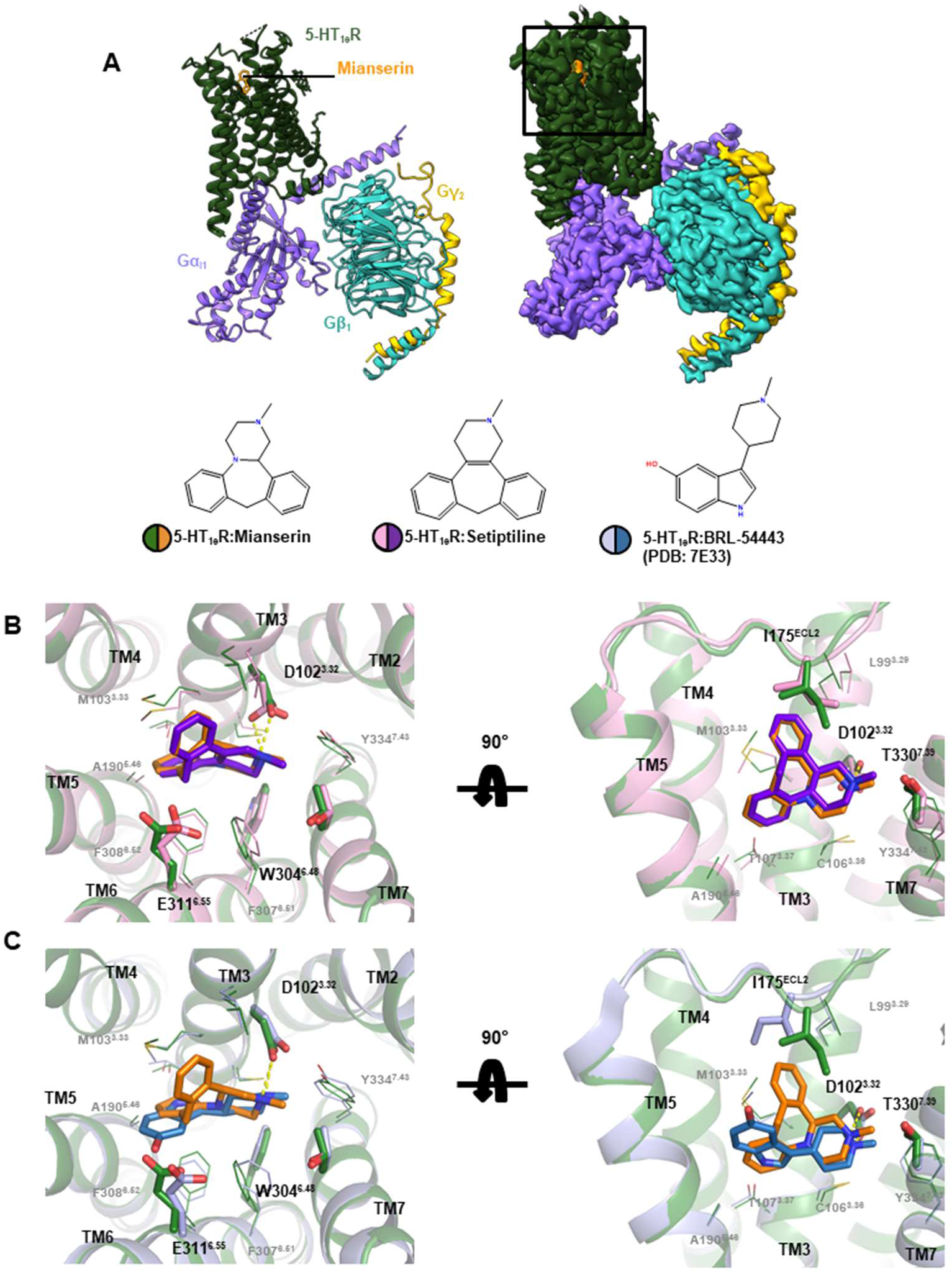
CryoEM structures of mianserin- and setiptiline-bound 5-HT_1e_R-Gα_i1_-Gβ_1_-Gγ_2_ complexes. **A,** 2D chemical structures of structurally characterized 5-HT_1e_R agonists (bottom) and overall structural model (left) and cryoEM map (right) of mianserin-bound 5-HT_1e_R-Gα_i1_-Gβ_1_-Gγ_2_ complex. Mianserin, 5-HT_1e_R, Gα_i1_, Gβ_1_, and Gγ_2_ are colored in orange, green, violet, teal, and yellow. **B**, superposition of orthosteric ligand binding pockets of mianserin- and setiptiline-bound 5-HT_1e_R. **C**, superposition of orthosteric ligand binding pockets of mianserin- and BRL-54443- bound 5-HT_1e_R. Key sidechains and drugs are shown as sticks, and a conserved ionic bond between drugs and D3.32 is shown as dashed yellow lines. Mianserin-5-HT_1e_R complex is colored in orange and green, setiptiline-5-HT_1e_R complex is colored in purple and pink, and BRL-54443-5-HT_1e_R complex (PDB: 7E33) is colored in dark blue and light blue.

Overall, our structures are similar to the previously published BRL-54443-bound 5-HT_1e_R- G_i1_ structure (*33*). Comparison of the mianserin- and setiptiline-bound complexes with that of the BRL-54443-bound complex yielded minimal global differences, with overall RMSDs of 0.66 Å and 0.63 Å, respectively (Fig. 3B-C, Fig. S5). The structures exhibit previously described hallmarks of activated GPCRs. For instance, the cytoplasmic tip of transmembrane helix (TM) 6 is moved outwards relative to inactive-state structures (discussed below) to accommodate the C-terminal alpha-helical domain of Gα_i1_. These changes likely originate from a torsion motion around F251^6.44^ of the P^5.50^-I^3.40^-F^6.44^ motif that allosterically links ligand- and transducer binding pockets (*36*). Compared to the previous structure (*33*), we observe additional densities for residues 158-160 at the extracellular side of TM4, the backbone of sidechains 170 and 171 at the end of extracellular loop (ECL) 2, and TM5 residues 213-217 (Fig. S5A, B). However, our structures lack density for the TM6 segment corresponding to residues 282-285. These minor differences do not appear to account for pharmacological differences between mianserin-/setiptiline- and BRL-54443-mediated activation, and likely reflect minor differences in sample and grid preparation. In contrast to the previous 5-HT_1e_R structure, we identify several densities corresponding to membrane components. For instance, we observe two cholesterols that are bound near the intracellular junction of TM2-4 (Fig. S5C). Two additional cholesterols are found on the interface formed by TMs 1 and 7, as well as TMs 1 and 2 (Fig. S5D). Lastly, a cholesterol appears to bind within the crevice formed by TM3-5, forming an interaction with the conserved Arg204^5.60^ (Fig. S5E).

### Structural, computational, and functional studies elucidate the unique binding pose of tetracyclic antidepressants at 5-HT_1e_R

Within the 5-HT_1e_R ligand binding pocket, mianserin and setiptiline exhibit similar topologies, adopting an open C-shaped bend across their tricyclic moieties and forming a conserved ionic bond between a tertiary amine in their fourth ring and D102^3.32^ (*37*) (Fig 3). The piperazine ring of mianserin (analogously the tetrahydropyridine ring of setiptiline) resides near the conserved residues F307^6.51^ and Y334^7.43^, which form one end of the orthosteric binding pocket. These interactions position the tricyclic moiety perpendicularly to the membrane between TM3, TM5, and TM6 (Fig. 3B). Both compounds appear to contact similar orthosteric binding pocket residues (within 4 Å), forming mostly hydrophobic interactions with L99^3.29^, M103^3.33^, C106^3.36^, T107^3.37^, I175^ECL2^, A190^5.46^, W304^6.48^, F308^6.52^, E311^6.55^, and T330^7.39^ (Fig. 3B).

The majority of binding pocket residues adopt similar conformations in the structures of TeCA-bound and BRL-54443-bound 5-HT_1e_R (*33*) (Fig. 3B-C). However, there were notable differences. The tricyclic moieties of mianserin and setiptiline assume a unique binding pose that places part of their scaffolds closer towards the extracellular space, where they occupy a site that has frequently been characterized as an extended binding pocket in other aminergic receptors (*38*). There, both drugs induce a displacement and rotation of I175^ECL2^ away from TM3 relative to BRL-54443 (Fig. 3C). In the setiptiline-bound structure, the electron density even suggests that I175^ECL2^ forms a more extended interface with the tricyclic moiety compared to that observed for mianserin (Fig. S3B, S4B). Furthermore, we observe a conformational change of E311^6.55^ in the mianserin-bound, but not the setiptiline-bound structure (Fig. 3B; Fig. S3, S4). To further investigate these subtle structural differences between mianserin-and setiptiline-bound 5-HT_1e_R binding pockets and validate our structural models, we conducted both computational and mutational studies.

### Molecular dynamics (MD) simulations and Structural Interaction Fingerprint (SIFt) analysis characterizes drug binding modes

To assess the stability of the binding modes of mianserin and setiptiline inferred by the density maps and uncover potential drug-related differences, we first performed four MD simulation replicas for a total of 1 µs for each ligand-receptor system and investigated ligand and receptor conformational flexibilities in an explicit lipid-water environment (see Methods section for details) (Fig. 4A-B). Both the receptor and ligands did not change significantly during the MD simulations as shown by their very low heavy-atom RMSD values (an average of 1.75 Å and 1.45 Å for mianserin and setiptiline, respectively, and an average of 1.69 Å and 1.49 Å for the receptor’s backbone in complex with mianserin and setiptiline, respectively (Fig. S6). A SIFt analysis (Fig. 4B) of the simulation trajectories confirmed that both mianserin and setiptiline maintain similar binding modes in the 5-HT_1e_R orthosteric binding pocket during simulations through similar interactions with receptor residues. Specifically, the tricyclic moiety in both mianserin and setiptiline interacted with probabilities larger than 60% with L99^3.29^, M103^3.33^, I175^ECL2^, A190^5.46^, F308^6.52^, and E311^6.55^ whereas the methylpiperazine group of both ligands formed apolar interactions with probabilities larger than 60% with C106^3.36^ and T330^7.39^ (Fig. 4B). The two ligands also formed a stable π-π stacking interaction with F308^6.52^ via the benzyl group of the tricyclic scaffold, as well as a hydrogen bond and an ionic bond with the side chain of D102^3.32^ via their piperazine nitrogen, which are typically present in aminergic GPCRs (*36, 37*). The most notable difference between the two ligands was an interaction formed with a probability larger than 30% by mianserin, but not setiptiline, with residue H177^ECL2^.

**Fig. 4.**
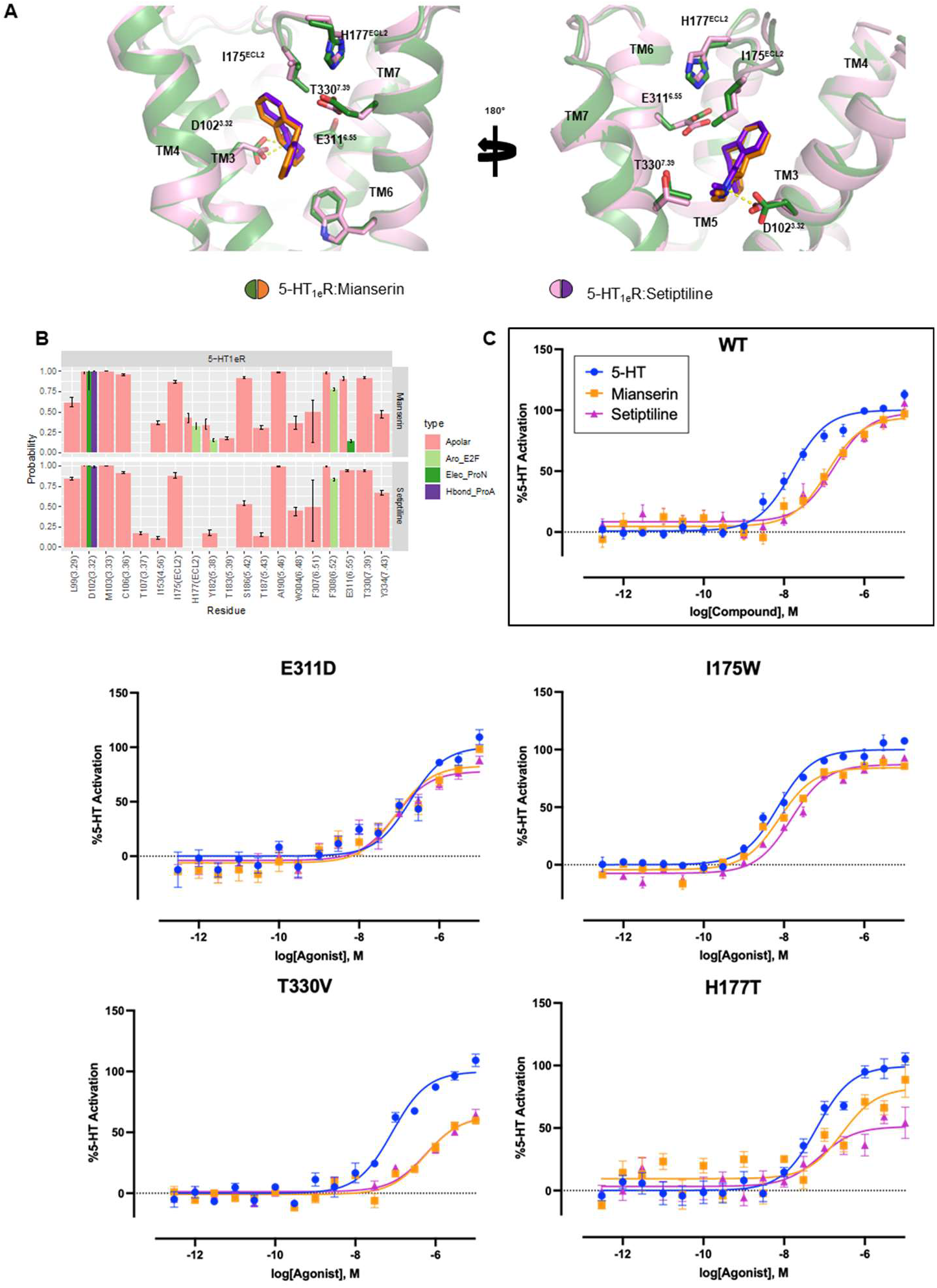
Experimental and computational structure-activity-relationship studies of mianserin and setiptiline interactions with 5-HT_1e_R binding pocket. **A,** views of the mianserin- and setiptiline-bound 5-HT_1e_R orthosteric pocket, with key residues interacting with drugs shown as sticks. Conserved ionic bond between drugs and D3.32 is shown as dashed yellow line. Mianserin-5-HT_1e_R complex is colored in orange and green, setiptiline-5-HT_1e_R complex is colored in purple and pink. **B**, structural interaction fingerprint (SIFt) analyses indicate probabilities of ligand−receptor interactions formed during four 250 ns MD simulations of the mianserin/setiptiline-5-HT_1e_R systems. The four interaction types formed by the ligands with the protein backbone and side chains are carbon−carbon atomic interactions (Apolar, pink), edge-to-face aromatic (Aro_E2F, light green), hydrogen bond with the protein as the hydrogen-bond acceptor (Hbond_ProA, purple), and electrostatic interaction with the protein negatively charged (Elec_ProN, light green). Only interactions with an average probability greater than 10% are displayed. **C**, concentration response experiments determining the activity of 5-HT, mianserin, and setiptiline at wildtype (wt) and mutant 5-HT_1e_R as measured by G_i1_ BRET in HEK293T cells. Experiments were performed in triplicate and mean±SEM from at least two (n=2) independent experiments were averaged and normalized to 5-HT’s response.

### Probing distinct 5-HT_1e_R binding pose of tetracyclic antidepressants via mutational analysis

Guided by our structural and computational analyses, we next interrogated the contributions of pocket residues to the distinct binding poses of mianserin and setiptiline when compared to the 5-HT analog BRL-54443. Towards this end we determined the activities of 5-HT, mianserin, and setiptiline in BRET G protein dissociation assays using mutant 5-HT_1e_R (Fig. 4C, Fig. S7, Table S4). Mutation of most ligand binding pocket residues non-specifically reduced the potency of agonist-mediated G protein dissociation. Interestingly, mutation of I175^ECL2^ to alanine reduced the potency of 5-HT relative to mianserin and setiptiline, while mutation to a bulkier phenylalanine did not impact potencies, and mutation to tryptophan rendered all three ligands roughly equipotent (Fig. 4C, Fig. S7, Table S4). This suggests that unlike the multicyclic compounds, potent receptor activation by 5-HT is reliant on the steric bulk near the extracellular boundary of the pocket, further highlighting the distinct receptor engagement of the antidepressants. Mutation of T330^7.39^ to a valine, as found in 5-HT_2_ receptors, reduced the efficacy of both mianserin (Emax= 64.4% of 5-HT) and setiptiline (Emax = 62.1% of 5-HT) (Fig. 4C, Fig. S7, Table S4). Similarly, H177^ECL2^, located next to E311^6.55^, when mutated to phenylalanine or alanine, also reduces mianserin’s efficacy (Emax= 60.5% and 62.2% of 5-HT, respectively). Notably, replacement with the polar residue threonine appeared to differentially impact the efficacies of setiptiline and mianserin (Emax = 51.2% and 78.3% of 5-HT, respectively), in line with the suggestion of different ligand-receptor interactions from our SIFt analysis.

E311^6.55^ shows different conformations in the mianserin- and setiptiline-bound structures and previous studies have highlighted the importance of interactions with residues at position 6.55 for ligand specificity (*39*). According to the SIFt analysis, E311^6.55^ forms highly probable apolar interactions with both mianserin and setiptiline during simulation, but it forms an electrostatic interaction with mianserin only, albeit with a very low probability (Fig. 4B). We thus probed the role of E311^6.55^ in ligand binding and/or receptor activation through substitution with glutamine, asparagine, or aspartate. We note that all mutations reduced ligands’ potencies to varying degrees. For instance, ablating the charge through an E311^6.55^Q mutation disproportionately reduced the potency of setiptiline relative to mianserin and 5-HT (Fig. S7, Table S4). Reducing the size through an E311^6.55^D mutation reduced potency disproportionately for 5-HT, rendering it equipotent to mianserin and setiptiline (Fig. 4C, Table S4). Reducing the size and removing the charge through an E311^6.55^N mutation resulted in a similar potency for 5-HT and setiptiline, with mianserin being marginally more potent (Fig. S7). This suggests that despite their high chemical similarity, there are subtle mechanistic differences in how mianserin and setiptiline bind to and activate 5-HT_1e_R, supporting our findings from the SIFt analysis, as well as the Shannon transfer entropy analysis reported below.

Together our experiments uncover and characterize how receptor residues differently contribute to the activity of the tetracyclic antidepressants compared to 5-HT. These data further suggest that rather than any individual interaction, mianserin’s and setiptiline’s overall binding pose, including interactions with extended pocket residues, is likely responsible for their unexpected agonism at 5-HT_1e_R (and likely 5-HT_1F_R).

### Molecular insights into the agonist activity of mianserin and setiptiline at 5-HT_1e_R

In the absence of any unambiguous drug-receptor interactions responsible for the agonist efficacy of the tested TeCA medications at 5-HT_1e_R, we turned to other approaches to analyze why these ligands are not antagonists, as with other 5-HTRs. We first compared our active state structures to the inactive state structures of 5-HTRs bound to chemically related antagonists. Specifically, we used inactive state structures of 5-HT_1B_R bound to the multicyclic research chemical methiothepin, and of 5-HT_2A_R bound to the multicyclic antipsychotic zotepine for comparison (*40, 41*). For ease of visualization, we show our analysis against only the mianserin- bound structure (Fig. 5A). As is the case for mianserin, both methiothepin and zotepine are anchored to their respective 5-HTRs via a conserved ionic bond with D^3.32^ (*37*), and their multicyclic moieties adopt a kinked C-shaped conformation occupying the amphipathic orthosteric binding pocket. However, the binding pose of both antagonists is fundamentally different, with both methiothepin and zotepine binding in a mirrored orientation compared to mianserin. As a result, the tricyclic pharmacophores moieties of the antagonists are oriented closer towards TM6, which shows distinct positioning in active- and inactive state structures (Fig. 5A). Since large scale TM6 movements concomitant with conformational changes in the rotameric toggle switch W^6.48^ are key elements for the transition from inactive to active state (*42, 43*), differences in TM6 interactions could conceivably contribute to the divergent pharmacology of the different multicyclic compounds.

**Fig. 5.**
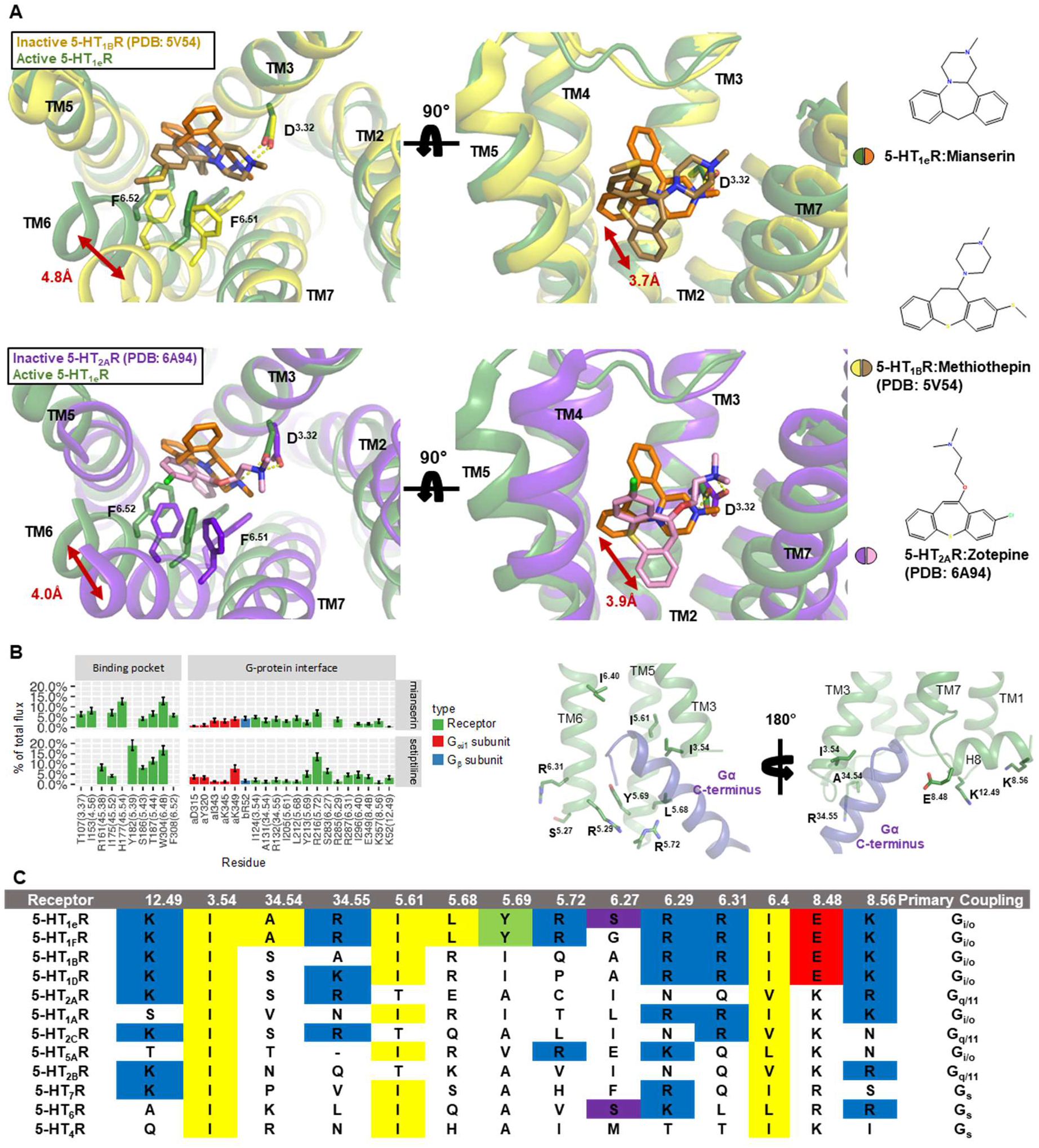
Molecular basis of mianserin’s and setiptiline’s agonism at 5-HT_1e_R. **A,** superposition of mianserin-bound 5-HT_1e_R and structures of inactive state 5-HTRs bound to multicyclic antagonists reveals key differences in drug binding poses. Mianserin-5-HT_1e_R complex is colored in orange and green, methiothepin-5-HT_1B_R complex (PDB: 5V54) is colored in brown and yellow, and zotepine-5-HT_2A_R complex (PDB: 6A94) is colored in pink and purple. Conserved ionic bonds between drugs and D3.32 are indicated as dashed yellow lines. Distances between mianserin and respective antagonists are measured between atoms closest to the 7TM core and shown as red arrows. 2D chemical structures of mianserin, methiothepin, and zotepine are shown on the right. **B**, Shannon transfer entropy analysis between pairs of ligand-residue or residue-residue contacts within a minimum distance of less than 4.5 Å during molecular dynamics simulations of mianserin-5-HT_1e_R-G_i1_ and setiptiline-5-HT_1e_R-G_i1_ complexes. Only residues that contribute significantly (>3%) to the transfer entropy between the ligand-binding pocket and the receptor G protein interface in each system are reported**. C**, sequence alignment of key residues of 5-HT receptor family involved in signal transduction.

The multicyclic pharmacophores also extend deeper into their respective binding pockets than mianserin. In fact, methiothepin and zotepine are bound 3.7 Å and 3.9 Å closer to the 7TM core compared to mianserin’s much shallower binding pose at 5-HT_1e_R (Fig. 5A). We further observe that comparison with inactive state structures of 5-HTRs bound to the chemically unrelated antagonist medications risperidone, aripiprazole, cariprazine, and ritanserin, similarly feature deep protrusions of the antagonists into the binding pocket relative to mianserin (Fig. S8). This further suggests that mianserin’s shallow binding pose might facilitate activation-related conformational changes near the 7TM core of the receptor that are otherwise inhibited by the deeper binding pose of antagonists.

Since allosteric communication between ligand and G protein binding sites plays a key role in the divergent pharmacological activities of drugs at GPCRs (*44*), we wanted to understand whether there was a discernable mechanistic basis in downstream allosteric networks by which mianserin and setiptiline induced agonism of 5-HT_1e_R. Towards this end we analyzed the contributions of residues to the allosteric communication between the orthosteric ligand binding site and the receptor-G protein interface via transfer entropy analysis of our MD simulations (>3% flux) (Fig. 5B). Notably, most of these contributing residues were the same for the two simulated drug−bound 5-HT_1e_R-G_i1_ systems (Fig. 5B), suggesting an overall common allosteric mechanism induced by the two drugs throughout 5-HT_1e_R. However, we do note differences for S238^6.27^ and R287^6.31^ at the receptor-G protein interface, as well as T107^3.37^, I153^4.56^, H177^ECL2^, Y182^5.39^, and F308^6.52^ in the binding pocket. Interestingly, some of the ligand-receptor interactions identified by the SIFt analysis (specifically, with residues C106^3.36^, E311^6.55^, T330^7.39^, and Y334^7.43^) do not appear to be involved in the allosteric coupling between the ligand binding pocket and the intracellular side of the receptor, suggesting that these interactions contribute to the binding of the ligand but not to the allosterically-regulated activation of the receptor. Notably, most of the residues establishing direct interactions with the ligands are conserved within the 5-HT_1_R family, except H177^ECL2^ (S190 in 5-HT_1A_, T203 and T192 in 5-HT_1B_R and 5-HT_1D_R, respectively). By contrast, we find that several of the residues contributing to the allosteric communication between the ligand binding pocket and the receptor-G protein interface are different among receptor subtypes (e.g., R132^34.55^ is N146 in 5-HT_1A_R, A159 in 5-HT_1B_R and K148 in 5-HT_1D_R; Y213^5.69^ is I266, I239 and I228 in 5-HT_1A_R, 5-HT_1B_R and 5-HT_1D_R, respectively; R216^5.72^ is T229 in 5-HT_1A_R, E242 in 5-HT_1B_R and P231 in 5-HT_1D_R) (Fig. 5C). This suggests that receptor-specific differences in allosteric communication could further contribute to differences in receptor subtype signaling in the 5-HT_1_R family, and thus help explain the unexpected agonism of mianserin and setiptiline at 5-HT_1e_R and 5-HT_1F_R.

## Discussion

Herein we report on the agonistic activity of multiple clinically utilized drugs, including those of the antidepressants mianserin, setiptiline, and mirtazapine, at the understudied serotonin receptors 5-HT_1e_R and 5-HT_1F_R. The finding that similar binding affinities across related cohorts of receptors may not translate to similar pharmacological activities has also previously been noted for other drugs such as LSD, which activates most serotonin receptors but is an antagonist at 5-HT_7_R (*45*). Our data thus further demonstrate that even within classes of GPCRs tuned to the same endogenous ligand, exogenous drugs have complex pharmacological profiles that cannot be generalized across receptor subtypes without the appropriate testing.

To investigate the binding modes and activation mechanisms of mianserin and setiptiline, we further determined cryoEM structures of drug-bound 5-HT_1e_R signaling complexes and combined our structural analysis with MD simulations and SAR studies. Accordingly, our studies suggest that a distinct binding mode together with receptor-specific allosteric communication between ligand and G protein binding sites is largely responsible for the unexpected agonism of mianserin and setiptiline at 5-HT_1e_R (and 5-HT_1F_R). Since none of our tested mutations convert mianserin or setiptiline into 5-HT_1e_R antagonists, we further support the notion that the overall ligand-binding pose and downstream allosteric networks, rather than solely specific ligand-receptor interactions, drive drug efficacy, as has been observed for opioid receptors and the β2-adrenergic receptor (*46, 47*).

The current degree of understanding about what differentiates an antagonist from an agonist at the structural level is relatively underdeveloped, and studies have shown that structural differences in the receptor binding pocket of active- and inactive state receptors can range from very subtle to dramatically different in a highly ligand-specific manner that is difficult to generalize, such as the 5-HT_2A_R agonists LSD and 25CN-NBOH compared to the antagonist methiothepin (*48*). However, in line with the herein reported findings, we have previously already shown that ergolines that penetrate deeper into the binding pocket of 5-HT_2B_R confer antagonism relative to chemically similar counterparts that exhibit a shallower binding mode (*49*). Structures of other drug-bound serotonin receptors similarly highlight how deep protrusions into the receptor core and distinct interactions with key activation motifs are a key feature of antagonists (*40, 41, 50, 51*). In addition, mianserin, setiptiline, and mirtazapine possess compact multicyclic scaffolds that span only 2-3 atoms between the anchoring amine and the aromatic rings. This likely facilitates contraction of the binding pocket, which is a key feature of GPCR agonism across several receptor classes (*42, 52, 53*). However, we also find that even larger multicyclic compounds such as chlorpromazine (Thorazine) can activate 5-HT_1e_R and 5-HT_1F_R, albeit with weaker potency compared to mianserin. Together, these findings suggest that the pockets of 5-HT_1e_R and 5-HT_1F_R allow for distinct positioning of these scaffold classes that facilitates contraction of the binding pocket and rearrangements in key activation motifs. A conclusive dissection of the mechanisms by which mianserin and setiptiline activate 5-HT_1e_R and 5-HT_1F_R, however, will require additional work including the characterization of the receptor’s inactive states, and/or structural studies of mianserin and setiptiline bound to 5-HTRs they antagonize.

Beyond providing much needed molecular and pharmacological insight into the mechanisms of 5-HT_1e_R and 5-HT_1F_R, our studies further have direct clinical implications. For instance, mianserin and mirtazapine have both been observed clinically to relieve migraines, and mirtazapine is currently even being used off-label for anti-migraine prophylaxis. We thus posit that the drugs’ potent agonism of the validated anti-migraine target 5-HT_1F_R contributes to their clinical anti-migraine efficacy (*31, 32*). Similarly, pimethixene showed initial promise as a treatment for migraine, albeit with a poor side effect profile attributed to its high affinity for H_1_R histamine receptors and 5-HT_2C_R serotonin receptors (*21, 22*). Lastly, methylergonovine has also shown clinical utility in treating migraine, although its long-term administration is contraindicated due to the potential for fibrotic symptoms as a consequence of 5-HT_2B_R agonism (*54*).

As for 5-HT_1e_R, little is known about its (patho)physiological role, outside of recent studies suggesting the receptor could be a potential drug target to treat ovarian cancer (*11*). However, 5-HT_1e_R is expressed across many brain areas including the basal ganglia, anterior cingulate cortex, and prefrontal cortex. These are key areas for the antidepressant action of increased serotonergic tone, and 5-HT_1e_R-mediated effects could thus conceivably contribute to the clinical efficacy of the antidepressants mianserin, setiptiline and mirtazapine. Furthermore, 5-HT_1e_R’s expression in the ovaries and uterus could explain the development of menstruation disorders in response to combining mirtazapine with other selective serotonin reuptake inhibitors (SSRIs) (*55*), or contribute to the clinical efficacy of methylergonovine as a uterotonic (*56*). These observations warrant further examination of the role of 5-HT_1e_R both neurologically and *in utero*, as well as the clinical effects of the herein described drugs that may, in part, be mediated by 5-HT_1e_R and/or 5-HT_1F_R.

Together, our studies not only uncover fundamental details about the unique molecular mechanisms of the enigmatic serotonin receptors 5-HT_1e_R and 5-HT_1F_R, but also suggest their direct involvement in the physiological effects of various drugs. Given the poorly understood physiological role of 5-HT_1e_R, we further hope to revitalize medicinal chemistry campaigns oriented around generating selective probes for future *in vivo* studies.

## Materials and Methods

### Construct Design for Protein Expression

Full-length human 5-HT_1e_R was cloned into a pFastBac vector similarly to the previously published structure of the receptor, featuring an N-terminal FLAG and 8x His tags followed by a TEV protease cleavage sequence and an N-terminal b562RIL apocytochrome (BRIL) fusion to improve expression and stability (*33*). Additionally, an L111^3.41^W mutation was added to stabilize 5-HT_1e_R (*57*). To simplify G protein heterotrimer expression and purification, we used a human Gγ_2_-Gα_i1_ fusion construct with a 3x GSA linker (gift of Dr. Tao Che) for formation of heterotrimeric G_i1_. The setiptiline–5-HT_1e_R–G_i1_ complex was further stabilized with a separately expressed ScFv16 (*58*). ScFv16 was cloned into a pFastBac vector with an N-terminal gp67 sequence to facilitate its secretion for subsequent purification (*59*). The mianserin–5-HT_1e_R–G_i1_ complex features a dominant-negative (DNGαi1) construct of Gαi1 containing the mutations S47N, G203A, E245A, and A326S (*60*).

### Protein Expression

5-HT_1e_R, Gγ_2_-Gα_i1_, and Gβ_1_ were expressed in *Spodoptera frugiperda* (*Sf9*) insect cells (Expression Systems) using baculoviruses. For the mianserin–5-HT_1e_R–G_i1_ complex, receptor and heterotrimeric G protein were expressed and purified separately, and assembled subsequently. For the setiptiline–5-HT_1e_R–G_i1_ complex, receptor and G protein components were co-expressed, and scFv16 was added after complex purification to stabilize the signaling complex. Cell cultures were grown in ESF 921 serum-free medium (Expression Systems) to a density of 2–3 million cells per ml and then infected with separate baculoviruses either at a multiplicity of infection of 3 (receptor alone) and 2:2 (Gγ2-Gαi1:Gβ1) in the case of the mianserin complex, or at a ratio of 2:1:1 (5-HT_1e_R:Gγ2-Gαi1:Gβ1) for the setiptiline complex. The culture was collected by centrifugation 48 h after infection and cell pellets were stored at −80 °C until further use.

### Protein Purification and Complexation

#### 5-HT_1e_R purification

5-HT_1e_R-expressing insect cells were disrupted by dounce homogenization in a hypotonic buffer containing 10 mM HEPES pH 7.5, 10 mM MgCl_2_, 20 mM KCl and home-made protease inhibitor cocktail (500 µM AEBSF, 1 µM E-64, 1 µM Leupeptin, 150 nM Aprotinin), and membranes were recovered as a pellet following centrifugation at 50,000 x G. Pelleted cellular membranes were homogenized and centrifuged twice in a high osmotic buffer containing 1 M NaCl, 10 mM HEPES pH 7.4, 10 mM MgCl_2_, 20 mM KCl and home-made protease inhibitor cocktail. Purified membranes were subsequently resuspended in a buffer of 150 mM NaCl, 10 mM MgCl_2_, 20 mM KCl, 20 mM HEPES pH 7.4, 10 uM mianserin, and home-made protease inhibitor cocktail, and agitated at room temperature for 30 minutes to allow compound binding, before being moved to 4 °C. Solubilization was initiated with the addition of Solubilization Buffer containing 150 mM NaCl, 20 mM HEPES pH 7.4, 2% n-Dodecyl-β-D-Maltopyranoside (DDM, Anatrace), and 0.4% CHS (Cholesteryl Hemisuccinate, Anatrace), and allowed to proceed for 2 hours with agitation. Subsequently, insoluble membrane components were removed from solution by centrifugation at 50,000 x G, and supernatant was supplemented with 20 mM imidazole and incubated overnight with TALON Superflow cobalt affinity resin (Cytiva). TALON resin was washed with Buffer 1 (800 mM NaCl, 20 mM HEPES pH 7.4, 0.1% DDM, 0.02% CHS, 10% glycerol, 20 mM imidazole and 10 µM mianserin) and Buffer 2 (800 mM NaCl, 20 mM HEPES pH 7.4, 0.05% DDM, 0.01% CHS, 10% glycerol, and 10 µM mianserin), and receptor was eluted with Buffer 3 (Buffer 2 with 250 mM imidazole). Eluted protein was concentrated in Vivaspin 6 Centrifugal Concentrators, desalted into a solution containing 100 mM NaCl, 20 mM HEPES pH 7.4, 0.05% DDM, 0.01% CHS, and 10% glycerol using PD MiniTrap Sample Preparation Columns, and allowed to equilibrate overnight after supplementation with 1% Lauryl Maltose Neopentyl Glycol (LMNG, Anatrace). LMNG-equilibrated samples were concentrated, flash frozen and stored at -80 °C for subsequent complexation.

#### G_i1_ purification

Gγ_2_-Gα_i1_/Gβ_1_-expressing insect cells were dounce homogenized in a lysis buffer consisting of 20 mM HEPES, pH 7.4, 100 mM NaCl, 1mM MgCl_2_, 10 µM guanosine diphosphate (GDP), 10% glycerol, 5 mM β-mercaptoethanol, 30 mM imidazole, 0.2% Triton X-100, and home-made protease inhibitor cocktail. The cytoplasmic and membrane fractions were separated by centrifugation at 50,000 x G for 20 min. The resulting supernatant was subjected to an additional centrifugation at 200,000 x G for 45 min to further clarify supernatant. The final supernatant was bound to HisPur Ni-NTA Resin (Thermo Scientific) overnight at 4 °C. Protein-bound Ni-NTA resin was washed with 20 cv of lysis buffer lacking 0.2% Triton X-100, followed by 20 cv lysis buffer lacking 0.2% Triton X-100 and 30 mM imidazole. Protein was eluted from the resin with lysis buffer lacking Triton X-100 and supplemented with 300 mM imidazole. Eluent from the first two elution fractions after the elimination of dead volume were concentrated using Vivaspin 6 Centrifugal Concentrators (Sartorius). Imidazole was removed from the concentrated eluent using PD MiniTrap Sample Preparation Columns (Cytiva) according to the manufacturer protocol. Eluted and desalted protein was injected onto a Superdex 200 Increase (Cytiva) size exclusion chromatography column equilibrated in 20 mM HEPES, pH 7.4, 100 mM NaCl, 1mM MgCl_2_, 0.01 mM guanosine diphosphate (GDP), 10% glycerol, 5 mM β-mercaptoethanol, and peak fractions containing intact heterotrimer were collected. Pooled fractions were concentrated, flash frozen and stored at -80 °C.

#### ScFv16 purification

ScFv16-expressing insect cells were pelleted by centrifugation, and the media they were grown in was collected for isolation of protein. Media was treated in sequence with Tris pH 8.0 (to a final concentration of 50 mM), CaCl_2_ (final concentration of 5 mM), and CoCl_2_ (final concentration of 1 mM), and stirred at room temperature for 1 hour to precipitate media components. Precipitate was allowed to sediment and further removed by filtration with a 0.22 μm PES Bottle Top Filter (Fisher). The final supernatant was supplemented with a final concentration of 10 mM imidazole and stirred with HisPur Ni-NTA Resin (Thermo Scientific) overnight at 4 °C. Protein-bound Ni-NTA resin was removed from supernatant by gradually removing solution from the top after sedimentation, and packed into a plastic flow column. Resin was subsequently washed with 10 column volumes (cv) of 20 mM HEPES pH 7.5, 500 mM NaCl, 10 mM imidazole, 10% glycerol. Further washing was done with 15 cv of 20 mM HEPES pH 7.5, 100 mM NaCl, 10% glycerol. Protein was eluted from the resin with 20 mM HEPES pH 7.5, 100 mM NaCl, 300 mM imidazole, 10% glycerol. The eluent was concentrated using Vivaspin 6 Centrifugal Concentrators (Sartorius). Imidazole was removed from the concentrated eluent using PD MiniTrap Sample Preparation Columns (Cytiva) according to the manufacturer protocol. Desalted protein was concentrated, flash frozen, and stored at -80 °C until needed for complexation.

#### Setiptiline/5-HT_1e_R–G_i1_ complex purification

For the setiptiline-bound 5-HT_1e_R–G_i1_ complex, cell pellets were thawed and resuspended in complex lysis buffer (75 mM NaCl, 10 mM MgCl_2_, 10 mM CaCl_2_, 20 mM KCl, 10 mM HEPES pH 7.4, 10 μM setiptiline, 10 μM GDP), dounce homogenized to lyse cells, and subsequently allowed to incubate for 30 minutes at room temperature with agitation. Apyrase (25 mU/mL, NEB) was then added, and the mixture was agitated for another 30 minutes at room temperature, when a 2-fold molar excess of ScFv16 (estimated from previous purification yields) was added and allowed to mix for another 30 minutes. The mixture was moved to 4 °C, and solubilization was initiated with the addition of Solubilization Buffer (described above) for 2 hours. Insoluble material was removed by centrifugation at 50,000 x G, and supernatant containing complex was supplemented with 20 mM imidazole, added to TALON Superflow cobalt affinity resin and agitated overnight at 4 °C. Resin was washed the following day with Complex Buffer 1 (Buffer 1 + 5 mM MgCl_2_), Complex Buffer 2 (Buffer 2 + 5 mM MgCl_2_), and eluted with Complex Buffer 3 (Buffer 3 + 5 mM MgCl_2_). Eluted complex fractions were pooled, concentrated with a Vivaspin 6 Centrifugal Concentrator, and supplemented with 1% LMNG, and allowed to sit for an hour at 4 °C. Protein was subsequently injected onto a Superdex 200 Increase (Cytiva) size exclusion chromatography column equilibrated in 20 mM HEPES, pH 7.4, 100 mM NaCl, 0.00075% LMNG, 0.00025% Glyco-diosgenin (GDN, Anatrace), 0.00015% CHS, 10 μM setiptiline and peak fractions containing intact receptor-heterotrimer-ScFv16 complex were collected. Pooled fractions were concentrated and kept on ice until CryoEM grid preparation.

#### Mianserin/5-HT_1e_R–G_i1_ complex assembly and purification

To form mianserin-bound 5-HT_1e_R-G_i1_ complexes, purified 5-HT_1e_R and G_i1_ were mixed on ice in a buffer with a final composition of 5 mM MgCl_2_, 10 mM CaCl_2_, 100 mM NaCl, 20 mM HEPES pH 7.4, 10 μM mianserin, and 10 μM GDP. Complexation was allowed to proceed for 30 minutes, after which apyrase was added and the complex mixture was incubated overnight at 4 °C. The following day, the complex mixture was injected onto a Superdex 200 Increase (Cytiva) size exclusion chromatography column equilibrated in 20 mM HEPES, pH 7.4, 100 mM NaCl, 0.00075% LMNG, 0.00025% Glyco-diosgenin (GDN, Anatrace), 0.00015% CHS, 10 μM setiptiline, and peak fractions containing intact receptor-heterotrimer complex were collected. Pooled fractions were concentrated and kept on ice until CryoEM grid preparation.

### CryoEM Grids and Imaging

Approximately 3 µl of 5-HT_1e_R–G_i1_ complex (at 4.5 and 9.8 mg/mL for mianserin- and setiptiline-bound complexes, respectively) sample was applied to glow-discharged Quantifoil R 1.2/1.3 300 mesh copper grids, which were subsequently blotted for 3 seconds and vitrified by plunging into liquid ethane using an EM-GP2 plunge-freezer (Leica). Grids were then stored in liquid nitrogen until data collection.

All automatic data collection was performed on a FEI Titan Krios instrument equipped with a Gatan K3 direct electron detector operated by the New York Structural Biology Center (New York, New York). The microscope was operated at 300 kV accelerating voltage, at a nominal magnification of 64,000x corresponding to a pixel sizes of 1.069 Å (mianserin-bound complex) and 1.076 Å (setiptiline-bound complex). For the mianserin-bound complex, 4,141 movies were obtained at a dose rate of 26.27 electrons per Å^2^ per second with a defocus ranging from −0.5 to −1.8 μm. The total exposure time was 2 s and intermediate frames were recorded in 0.05 s intervals, resulting in an accumulated dose of 52.54 electrons per Å^2^ and a total of 40 frames per micrograph. For the setiptiline-bound complex, 6,935 movies were obtained at a dose rate of 25.77 electrons per Å^2^ per second with a defocus ranging from −0.5 to −1.8 μm. The total exposure time was 2 s and intermediate frames were recorded in 0.05 s intervals, resulting in an accumulated dose of 51.53 electrons per Å^2^ and a total of 40 frames per micrograph.

Movies were motion-corrected using MotionCor2 (*61*) and imported into cryoSPARC (*62*) for further processing. CTFs were estimated using patchCTF in cryoSPARC. An initial model was produced from a subset of micrographs using blob picking, followed by extraction, 2D classification, selection of key classes, and generation of a model ab initio. Subsequent models were produced from curated micrograph sets using particles found by template picking using the initial model. Particles were extracted, subjected to 2D classification, and a final particle stack was obtained by iterative rounds of multi-class 3D heterogeneous refinement sorting with several bad densities from rejected particles and the best density from each round of classification. Finally, non-uniform refinement was used to further refine the receptor-G protein complexes to their final resolutions – a stack of 1,321,565 particles for the mianserin-bound complex at 3.30 Å, and a stack of 221,798 particles for the setiptiline-bound complex at 3.32 Å. In the case of the mianserin-bound complex, we found that further attempts to reduce the particle stack size reduced the global resolution substantially, whereas the setiptiline-bound complex readily split into a small fraction of the particle stack size with a similar resolution.

### Model Construction and Refinement

Model building was conducted in COOT (*63*), using the published 5-HT_1e_R-G_i1_ complex as a template. Manual adjustments were iteratively performed in alternation with real-space refinement using the real_space_refine PHENIX program (*64*) to obtain the final refined atomic model, which was validated using MolProbity (*65*). The final mianserin-and setiptiline-bound complex models include receptor residues 20 to 160, 170 to 216, and 283 to 358. Poorly resolved residues were modeled as alanines. Ligand models were generated with the Grade2 webserver (*66*). All structural figures in this text were rendered using PyMol (*67*), except for the overall density map in Figure 3, and local resolution figures in the supplement, which were made using ChimeraX (*68*).

### Mammalian Cell Culture

HEK239T cells and HTLA cells, a modified HEK293T line expressing a modified β-arrestin (a gift from Dr. Bryan Roth), were used for assays presented in this work, and cultured in Dulbecco’s Modified Eagle Medium (DMEM, Gibco) supplemented with 10% v/v Fetal Bovine Serum (FBS) and 1% v/v penicillin-streptomycin (P/S). Intermittently, HTLA cells were cultured with selection media that contained 100 μg/mL of hygromycin and 2 μg/ml of puromycin to maintain the stability of transgenic constructs in the cell line. All cell plates were maintained in a humid 37° incubator with 5% CO_2_.

### BRET G Protein Dissociation Assays

HEK293T cells were seeded in 10 cm plates or six-well plates, transferred into DMEM containing 1% v/v dialyzed FBS, and allowed to equilibrate for at least an hour at 37°C. Equilibrated cells were then transfected using DNA/PEI particles at a 1:2 ratio in OptiMEM (Gibco). For con-transfection we used a ratio of 1:3:3:3 of receptor:Gα:Gβ:Gγ using TRUPATH plasmids (*18*) as well as 5-HT_1e_R and 5-HT_1F_R cloned into pcDNA3.1 vectors. On the day following transfection, cells were plated in a white, non-transparent bottom 384-well plate at a density of 10,000 cells per well. The next day, media was exchanged in each well for 30 μl Assay Buffer (20 mM HEPES pH 7.4, 0.1% w/v BSA, 0.01% v/v ascorbic acid, 1x Hank’s Balanced Salt Solution (HBSS, Gibco)), and 15 μl of compounds diluted in series in Assay Buffer were added to the appropriate wells. Cells were incubated for 20 minutes in a 37°C incubator, and then brought to room temperature to incubate for 10 minutes. Subsequently, 15 μl of 30 μM coelenterazine 400a (GoldBio) in Assay Buffer was added to all wells, and the plate was immediately read using a multimode fluorescent plate reader (Perkin Elmer Victor NIVO). Emission filters were set to 395 nm (RLuc8-coelenterazine 400a) and 510 nm (GFP2) with integration times of 1 second per well. Data was plotted and analyzed using GraphPad Prism 8.0. Functional activity, the G Protein Dissociation, was determined by calculating the ratio of GFP2 to Rluc8 emission counts. These ratios were then plotted as a function of drug concentration and then analyzed in a non-linear regression analysis of log(agonist) versus response. Data was then normalized as a percentage of serotonin activation.

### cAMP Accumulation Assays

HEK293T cells were seeded in 10 cm plates, transferred into DMEM containing 1% v/v dialyzed FBS, and allowed to equilibrate for at least an hour at 37°C. Equilibrated cells were transfected with 0.8 μg of 5-HT_1e_R or 5-HT_1F_R and 8 μg of GloSensor plasmid (Promega), after mixing with 17.6 μl of PEI in OptiMEM (Gibco). On the day following transfection, cells were plated in white, transparent bottom 384-well assay plates at a density of 10,000 cells per well. The following day, media was exchanged in each well for 30 μL of Assay Buffer containing 1.2 mM of D-Luciferin (GoldBio). Cells were incubated for at least one hour at 37°C before the addition of 15 μl of compounds diluted in series in Assay Buffer. Cells were allowed to incubate in the dark at room temperature for 30 minutes. Subsequently, 15 μl of 1.6 μM isoproterenol in Assay Buffer was added to each well to stimulate cAMP production. Cells were incubated in the dark at room temperature for an additional 15 minutes before being read in a Perkin Elmer Trilux Microbeta. Luminescent counts per second (LCPS) were reported and then plotted as a function of drug concentration and analyzed in a non-linear regression analysis of log(agonist) versus response in GraphPad Prism 8.0.

### Arrestin Recruitment Assays

For arrestin recruitment we used the PRESTO-Tango assay, which was performed essentially as described (*14*). The 5-HT_1e_R and 5-HT_1F_R PRESTO-Tango constructs were obtained from Addgene. For transfection, 8 μg of DNA and 16 μl of PEI were incubated in OptiMEM for 20 min, and then added to HEK293T cells in DMEM supplemented with 1% v/v dialyzed FBS in 10 cm plates. After transfection, cells were placed into the 37°C incubator overnight. The following day, cells were plated in 40 µl DMEM in supplemented with 1% v/v dialyzed FBS in wells of poly-lysine coated white, transparent bottom 384-well plate as described above. The cell plates were placed into the 37°C incubator for approximately 4 hours, or until cells adhered to the bottom of the plate. The assay buffer used for the PRESTO-Tango assay was similar to the one used for the TRUPATH assay, but supplemented with 0.3% BSA and 0.03% ascorbic acid. 20 μl of drugs at 3x concentration in assay buffer were then added to the cells, and the plates were incubated at 37°C overnight for approximately 16 hours. The next day, the drug solution and media were removed and replaced with 20 μl of BrightGlo Reagent (Promega). Cells were incubated in the dark at room temperature for 20 minutes before being read in a Perkin Elmer Trilux Microbeta. Luminescent counts per second (LCPS) were reported and then plotted as a function of drug concentration and analyzed in a non-linear regression analysis of log(agonist) versus response in GraphPad Prism 8.0.

### Screening Analysis

The compound screen was done testing each compound in quadruplicate at a dose of 10 μM. The quadruplicates were averaged and the means were normalized against a baseline: Every compound that elicited a response 0.5*log2 fold change over baseline was considered active. For the GloSensor Assay, the baseline was set via a control plate, which had cells transfected only with GloSensor, while PRESTO-Tango assays did not use a control plate. For the GloSensor cAMP assay, compounds were considered active that showed a more than 0.5*log2 fold change in signal in receptor transfected cells compared to cells that were only transfected with the cAMP sensor. For PRESTO-Tango, compounds were considered active that showed a more than 0.5*log2 fold change in signal compared to DMSO. All concentration response experiments reported herein were performed in triplicate and averaged from at least 2 independent experiments (detailed biological replicate numbers are indicated in figure legends). Data is shown as mean±SEM All analyses were done in GraphPad Prism.

### System setup for MD simulations

The cryoEM structures of human 5-HT_1e_R in complex with Gα_i1_β_1_γ_2_ and bound to mianserin or setiptiline were used as starting points for MD simulations after adding atomic coordinates of the missing ECL2, which was built ab initio with RoseTTAFold (*69*). The N-terminal, C-terminal, and intracellular loop 3 (ICL3) regions of 5-HT_1e_R, which were missing from the two cryoEM structures, were not included in the modeling and the simulations. Instead, the truncated receptors were capped with acetyl- and N-methyl-groups at the N- and C-terminal, respectively. While the missing short Leu234-Met240 loop of Gα_i1_ was built *ab initio* with RoseTTAFold, the missing N-terminal and Ile55-Thr181 regions of Gα_i1_, as well as N- and C-terminal regions of the β and γ subunits of G_i_, were not included and their ends capped with acetyl- and N-methyl groups, respectively. The Protein Preparation Wizard tool (*70*) of Maestro v. 12.6 (Schrödinger) was used to assign bond orders and add cap terminal groups and missing hydrogens to both the receptor and G protein, as well as most probable protonation states according to a pH of 7.4. Following the PROPKA (*71*) hydrogen-bond network optimization protocol at pH 7.4, the protein systems underwent restrained energy minimization using the OPLS3e force field (*72*) and default parameters until the root mean square deviation (RMSD) of the protein heavy atoms reached a value of 0.3 Å.

The three-dimensional structure of S(+)-mianserin, was obtained from the Cambridge Crystallographic Data Centre (CCDC) website. S(+)-mianserin and setiptiline ligands with the most probable protonation states at pH of 7.4 were prepared using the default LigPrep protocol in the Schrödinger software release 2020-4 using OPLS3e partial charges (*72*). The dihedral parameters of S(+)-mianserin and setiptiline that were not included in the standard OPLS3e force field were generated using the Force Field Builder (FFBuilder) tool of the Schrödinger suite.

### Molecular dynamics simulations

Each ligand-bound receptor system was embedded in a 1-palmitoyl-2-oleoylphosphatidylcholine (POPC) bilayer, solvated in an orthorhombic box using the SPC water model (*73–75*) with a 10 Å distance in each cartesian direction between the protein and box boundaries, and neutralized with Na^+^ and Cl^-^ ions. A concentration of 0.15 M of NaCl was added to mimic physiological conditions. The OPLS3e force field was used for system preparation using the Desmond System Builder tool of Maestro (*76*).

MD simulations were run using Desmond’s default simulation parameters, including a 2 fs time step for bonded forces and short-range nonbonded forces and a 6 fs time step for long-range nonbonded forces using the reversible reference system propagation algorithm (RESPA) integrator. Following the standard membrane relaxation protocol in Desmond, constant-pressure and constant-temperature (NPT) equilibration runs were carried out in 9 steps, the first 8 of which employing gradually relaxing positional restraints on the heavy atoms of lipids, protein side chains, protein backbone, ligand ring atoms, and, lastly, the remaining ligand atoms. The last equilibration step consisted of a 1 ns unrestrained NPT run. During equilibration, the system temperature and pressure were maintained at 300 K and 1 bar, respectively, using the Nose-Hoover thermostat (*77*) and a semi-isotropic Martyna, Tobias, and Klein (MTK) barostat. Short-range Coulomb interactions were cut at 9 Å.

Four independent 250 ns MD production runs were carried out for each ligand−receptor complex, and structural data collected every 0.5 ns. The resulting 2000 frames for each system were stripped of the ions and lipids prior to aligning the heavy atoms of their respective energy-minimized 5 HT_1e_R cryoEM structures and to calculating their RMSD for both the receptor and the ligand’s heavy atoms, using an in-house python script.

### Structural Interaction Fingerprint (SIFt) Analyses

SIFt analyses (*78*) were performed on the merged MD simulation trajectories using an in-house python script. The interactions between ligands and receptor residues – both backbone and side chains – were calculated as a 9-bit representation based on (i) hydrogen-bond interactions with the protein as the hydrogen-bond donor (Hbond_proD) or hydrogen-bond acceptor (Hbond_proA), (ii) apolar interactions (carbon−carbon atoms in contact), (iii) face-to-face (Aro_F2F) and edge-to-face (Aro_E2F) aromatic interactions, (iv) electrostatic interactions with positively (Elec_ProP) or negatively charged (Elec_ProN) residues, and (v) one-water-mediated hydrogen bond (Hbond_1wat) or two-water-mediated hydrogen bonds (Hbond_2wat). A distance cutoff of 4.5 Å was used to define apolar interactions, a cutoff of 4 Å was used to describe aromatic and electrostatic interactions, and a cutoff of 3.5 Å was used to describe H-bond interactions. The probability of each interaction was estimated using a two-state Markov model, and sampling the transition matrix posterior distribution using standard Dirichlet priors for the transition probabilities as described (*79*).

### Transfer Entropy Analyses

To elucidate the information flow across each simulated ligand-bound 5-HT_1e_R-Gα_i1_β_1_γ_2_ system we calculated the Shannon transfer entropy (*80*) between pairs of ligand-residue or residue-residue contacts within a minimum distance of less than 4.5 Å during MD simulations. Specifically, transfer entropy values were obtained with the MDEntropy package (*81*) between 14 ligand−residue and 696 residue−residue contacts in the case of the mianserin-bound system, and 12 ligand−residue and 643 residue-residue contacts in the case of the setiptiline-bound system, yielding transfer entropy matrices of 710 and 655 structural descriptors, respectively. The NetworkX python library (*82*) was used to build a directed graph in which each node represented a direct ligand−residue or residue−residue contact and the “length” of the edge between two nodes corresponded to the negative logarithm of the transfer entropy between them. The information flow from the ligand orthosteric binding site to the receptor-G protein interface was described by the set of paths Γ connecting the set of ligand−residue contacts (source set) to the set of contacts between receptor and G-protein residues (target set). The flux *F_y_* through each path was calculated as the total length of the path, and the contribution *C_k_* of each node *k* to the allosteric communication between the ligand binding pocket and the receptor-G protein interface was measured by normalizing the flux and summing over all paths going through that node as:

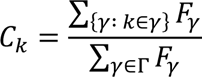

To assess the contribution of each residue to the allosteric mechanism, we calculated the sum of the contributions of all the nodes that contained that residue. Only residues that contributed the most to allosteric communication (>3% flux) were considered.

## Acknowledgments

We would like to acknowledge the support of the structural work by members of the National Center for cryo-EM Access and Training (NCCAT) and the Simons Electron Microscopy Center located at the New York Structural Biology Center, supported by the NIH Common Fund Transformative High Resolution Cryo-Electron Microscopy program (U24 GM129539,) and by grants from the Simons Foundation (SF349247) and NY State Assembly. We further acknowledge the staff at the Laboratory for BioMolecular Structure (LBMS), which is supported by the DOE Office of Biological and Environmental Research (KP160711). This work was supported in part through the computational resources and staff expertise provided by Scientific Computing at the Icahn School of Medicine at Mount Sinai and supported by the Clinical and Translational Science Awards (CTSA) grant UL1TR004419 from the National Center for Advancing Translational Sciences. We would also like to acknowledge Dr. John McCorvy for critical evaluation of the manuscript.

## Author contributions

G.Z., A.L.W., B.F., M.F., and D.W. conceptualized the study. G.Z. and A.L.W. performed the structural studies. G.Z., A.K.P., and A.L.W. performed the pharmacological assays. B.F. performed the MD simulations under the supervision of M.F. All authors contributed to data analysis and interpretation. G.Z. and D.W. wrote the manuscript with contributions of all authors.

## Data and materials availability

All data to evaluate the findings in this manuscript are available in the main text or the supplementary materials. Density maps and structure coordinates have been deposited in the Electron Microscopy Data Bank (EMDB) and the PDB: mianserin/5-HT_1e_R-Gα_i1_-Gβ_1_-Gγ_2_ (EMD-XXXXX and PDB XXX) and setiptiline/5-HT_1e_R-Gα_i1_-Gβ_1_-Gγ_2_/scFV16 (EMD-XXXXX and PDB XXX). Correspondence and requests for materials should be addressed to.

## Supplementary Materials

**Fig. S1.**
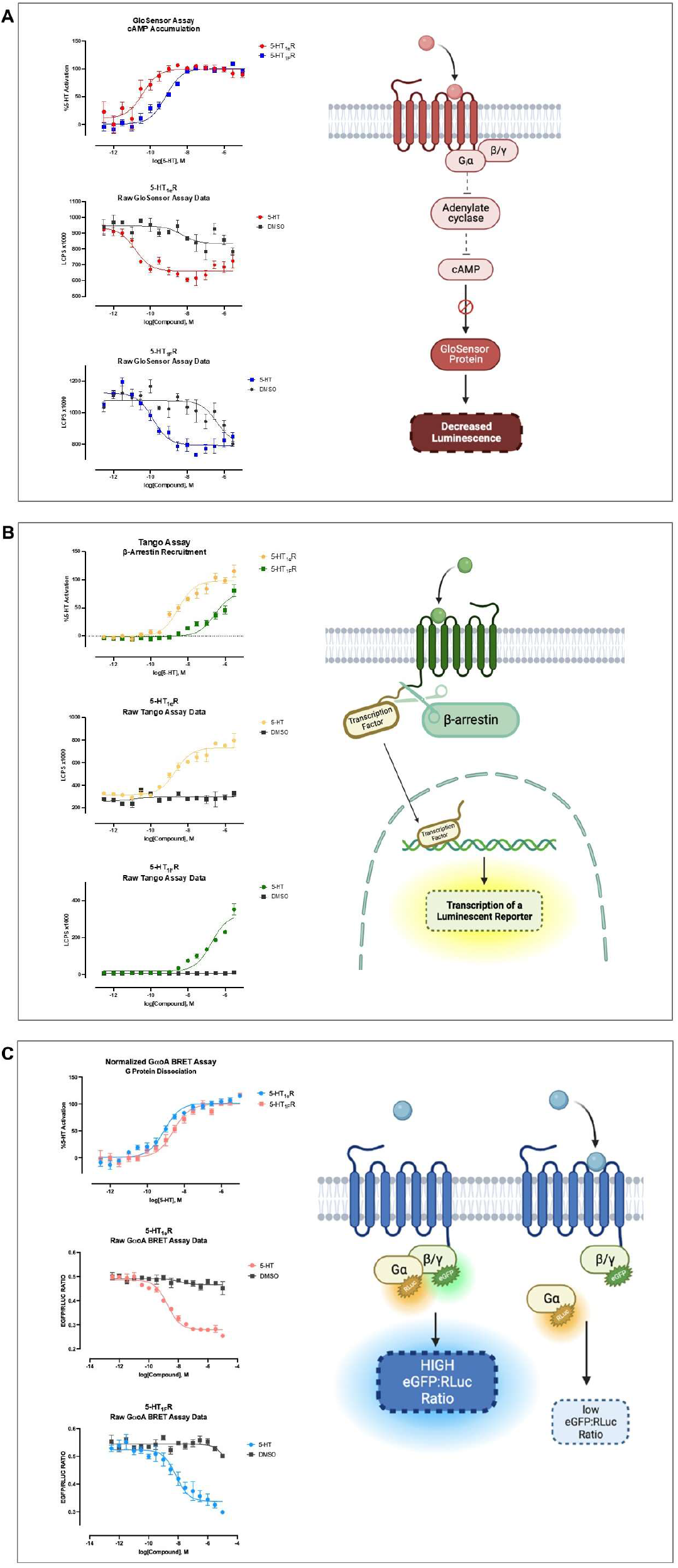
5-HT_1e_R and 5-HT_1F_R signaling assays. Schematics and example data are shown for GloSensor cAMP accumulations assay (**A**), TRUPATH bioluminescence resonance energy transfer (BRET) G protein activation/dissociation assay (**B**), and PRESTO-TANGO β-arrestin recruitment assay (**C**) used throughout this study. We further show data from concentration response experiments in HEK293T cells using either 5-HT or DMSO (negative control) either as normalized (% 5-HT response) or raw data. Raw data is shown as luminescence counts per second (LCPS) for cAMP and PRESTO-TANGO, and the ratio of eGFP fluorescence counts divided by RLuc bioluminescence counts for BRET. All experiments were performed in triplicate and mean±SEM from three (n=3) independent experiments were averaged and normalized data was calculated based on 5-HT’s response.

**Fig. S2.**
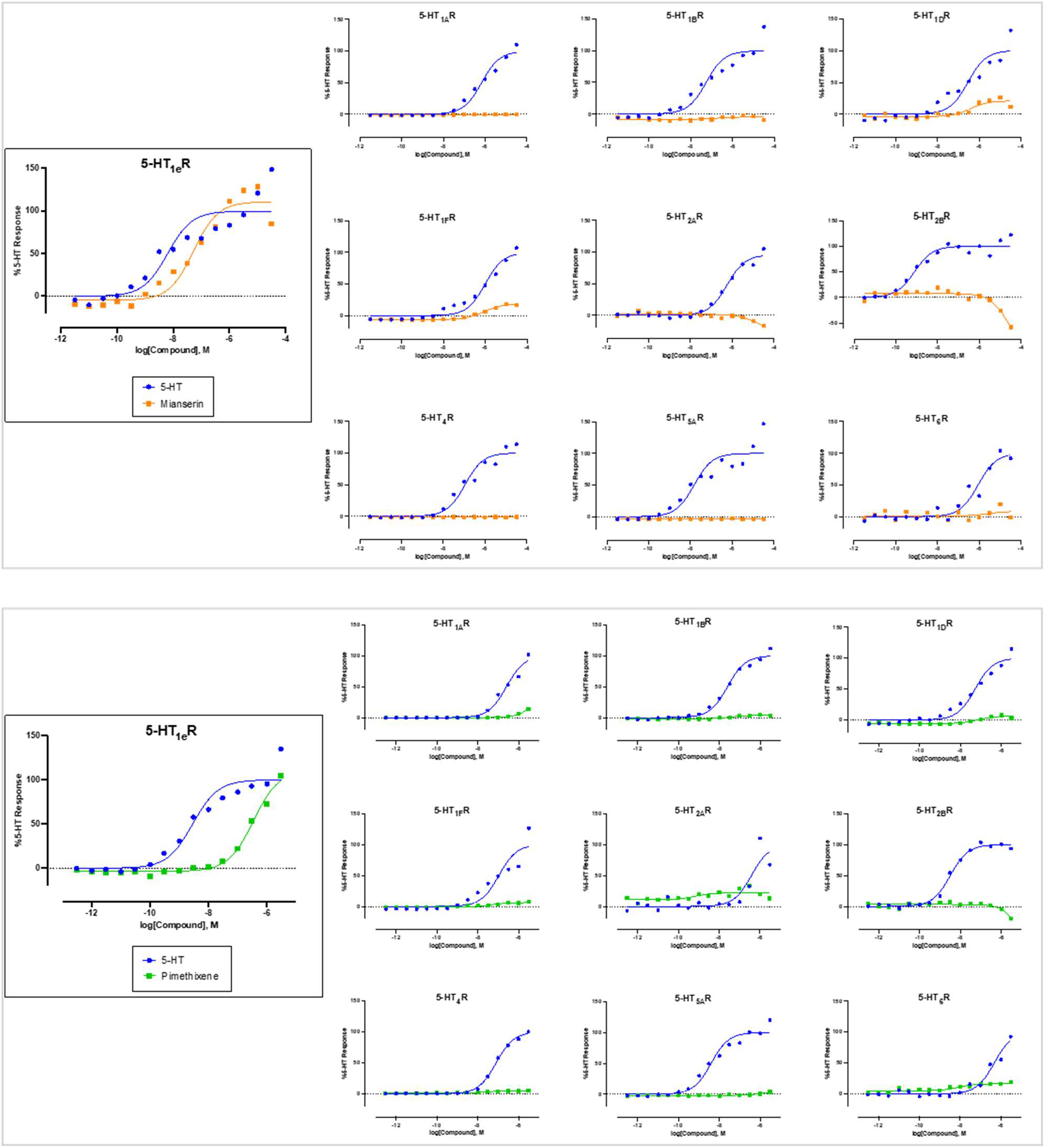
Activity of mianserin and pimethixene at 5-HT receptors. β-arrestin recruitment experiments in HEK293T cells were performed in triplicate. Mean±SEM from one (pimethixene, n=1) and two (mianserin, n=2) independent experiments were averaged and normalized to 5-HT’s response. Mianserin was a full and potent agonist at 5-HT_1e_R (pEC_50_ = 7.28 ± 0.10; Emax = 115.7 ± 4.8% of 5-HT), and a weaker partial agonist at 5-HT_1F_R (pEC_50_ = 6.07 ± 0.06; Emax = 24.5 ± 0.8% of 5-HT) and 5-HT_1D_R (pEC_50_ = 6.43 ± 0.24; Emax = 25.1 ± 2.8% of 5-HT). Pimethixene was a full, albeit less potent, agonist at 5-HT_1e_R (pEC_50_ = 6.46 ± 0.04; Emax = 114.4 ± 3.2% of 5-HT), but not active at either 5-HT_1F_R or 5-HT_1D_R.

**Fig. S3.**
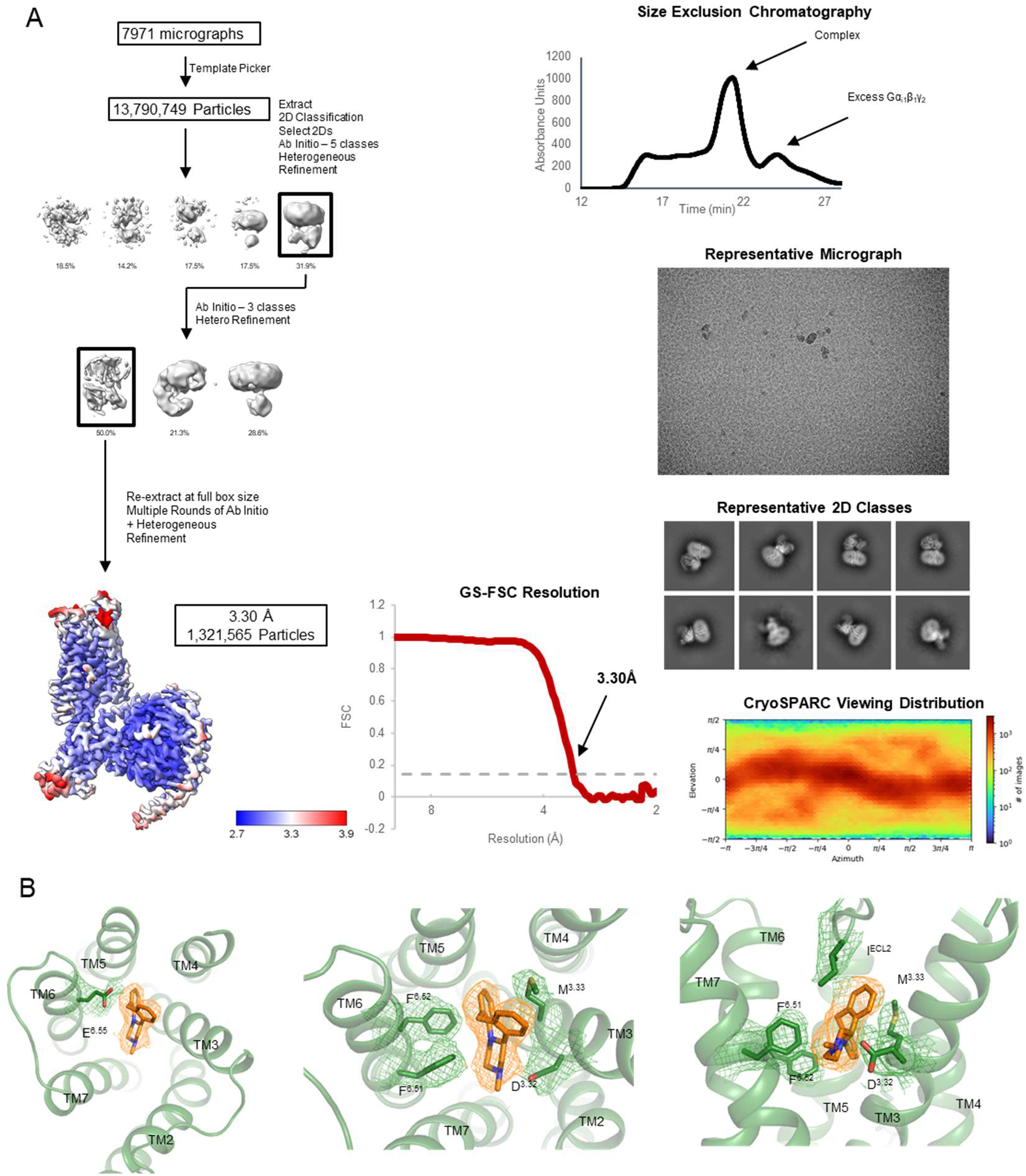
CryoEM structure determination of mianserin-bound 5-HT_1e_R-Gα_i1_-G_β1_-G_γ2_. **A**, Analytical size exclusion chromatography shows monodisperse and pure protein of drug-bound 5-HT_1e_R-G_i1_ complex. CryoEM data were collected on 300 keV Krios, a representative micrograph is shown, and processed in cryoSPARC: particles were picked from motion corrected micrographs, subjected to 2D classification (representative classes are shown), followed by ab initio model building and 3D classification. After multiple rounds of 3D classification, the final particle stack was subjected to local CTF refinement followed by local refinement. A final map was obtained with GS-FSC indicating a global resolution of 3.30 Å applying the 0.143 cutoff. Viewing direction distribution analysis (cryoSPARC) indicates isotropic distribution of views and sufficient coverage. An initial model was built in PHENIX, and then further refined in ServalCat for the generation of final maps and coordinates. Calculations in cryoSPARC indicate local resolutions of up to 3 Å around orthosteric binding site. **B**, Sharpened cryoEM density of mianserin (orange) and interacting residues (green) at a contour level of 5σ.

**Fig. S4.**
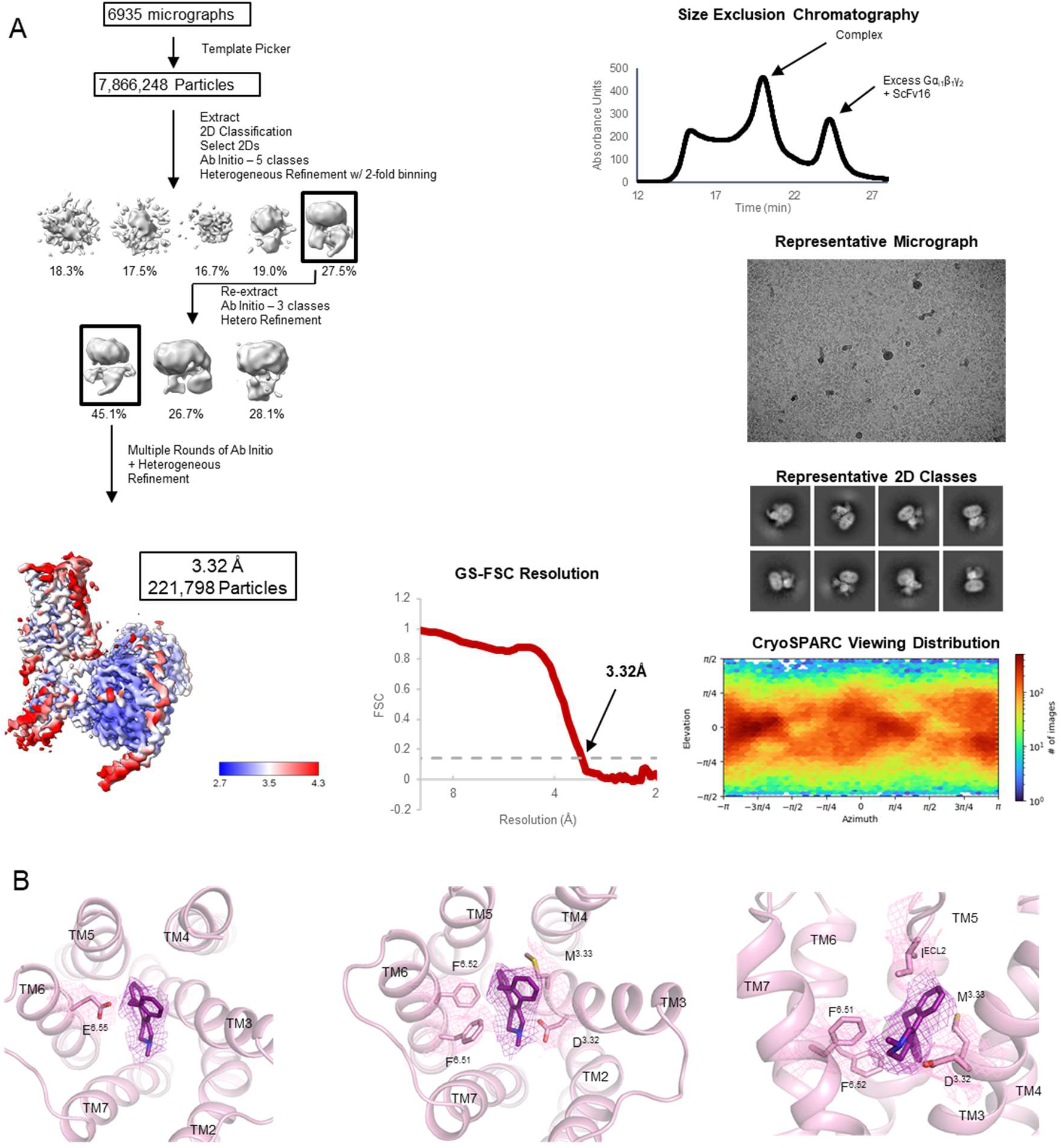
Cryo-EM structure determination of setiptiline-bound 5-HT_1e_R-Gα_i1_-G_β1_-G_γ2_– scFV16. **A**, Analytical size exclusion chromatography shows monodisperse and pure protein of drug-bound 5-HT_1e_R-G_i1_ complex. CryoEM data were collected on 300 keV Krios, a representative micrograph is shown, and processed in cryoSPARC: particles were picked from motion corrected micrographs, subjected to 2D classification (representative classes are shown), followed by ab initio model building and 3D classification. After multiple rounds of 3D classification, the final particle stack was subjected to local CTF refinement followed by local refinement. A final map was obtained with GS-FSC indicating a global resolution of 3.32 Å applying the 0.143 cutoff. Viewing direction distribution analysis (cryoSPARC) indicates isotropic distribution of views and sufficient coverage. An initial model was built in PHENIX, and then further refined in ServalCat for the generation of final maps and coordinates. Calculations in cryoSPARC indicate local resolutions of up to 3 Å around orthosteric binding site. **B**, Sharpened cryoEM density of setiptiline (purple) and interacting residues (pink) at a contour level of 5σ.

**Fig. S5.**
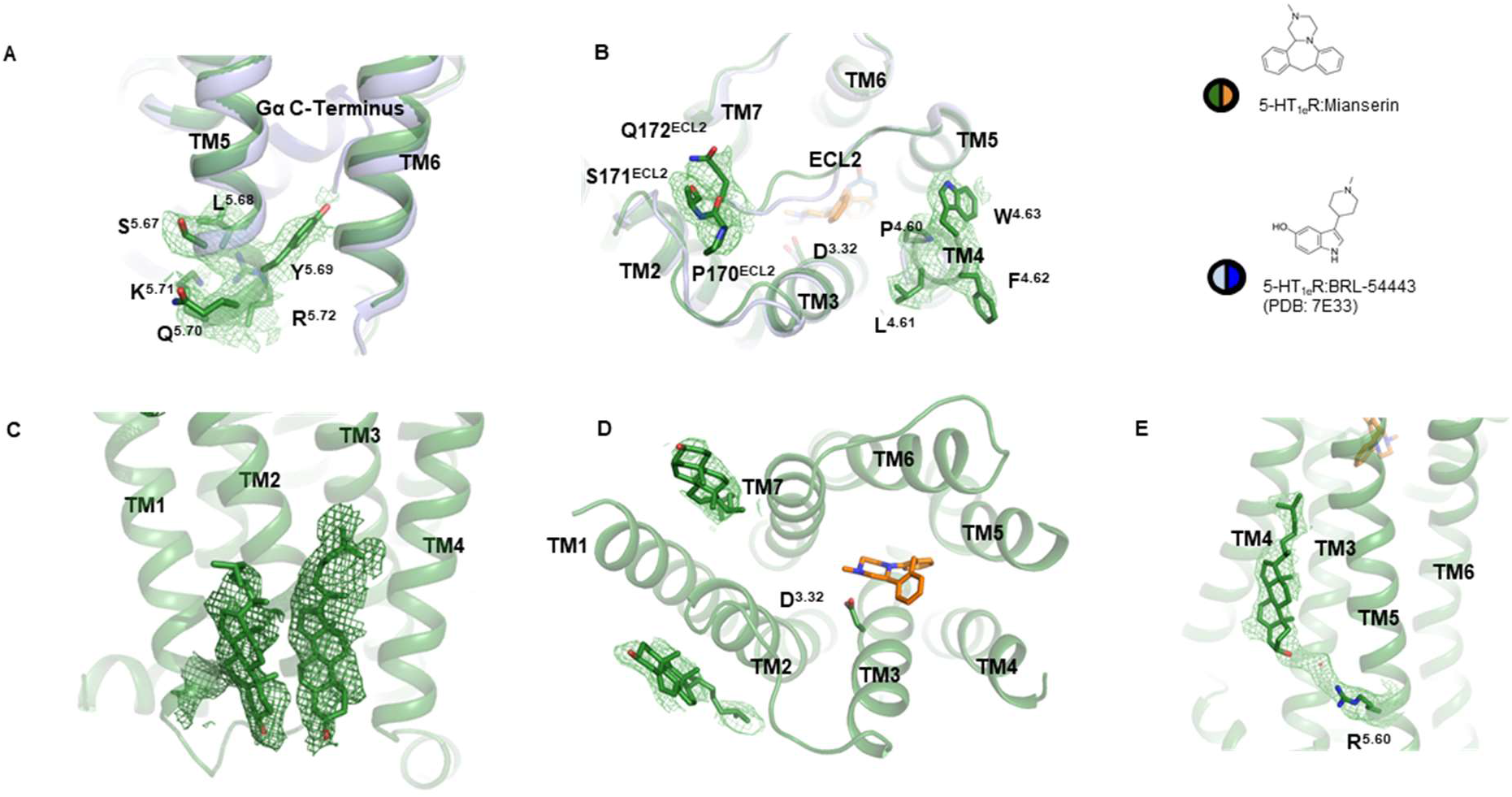
Differences to structure of BRL-54443-bound 5-HT_1e_R-G_i1_ complex (PDB: 7E33). Densities and model of the herein reported mianserin-bound 5-HT_1e_R-Gα_i1_-Gβ_1_-Gγ_2_ complex are shown in green and compared to the previously reported BRL-54443-bound 5-HT_1e_R-G_i1_ complex (PDB: 7E33) shown in light blue. Our map allowed to model additional receptor residues near the cytoplasmic tip of TM5 (**A**) and the extracellular surface (**B**). We also observed density for multiple receptor-bound sterols (**C-E**). All cryoEM densities are shown at a contour level of 5σ.

**Fig. S6.**
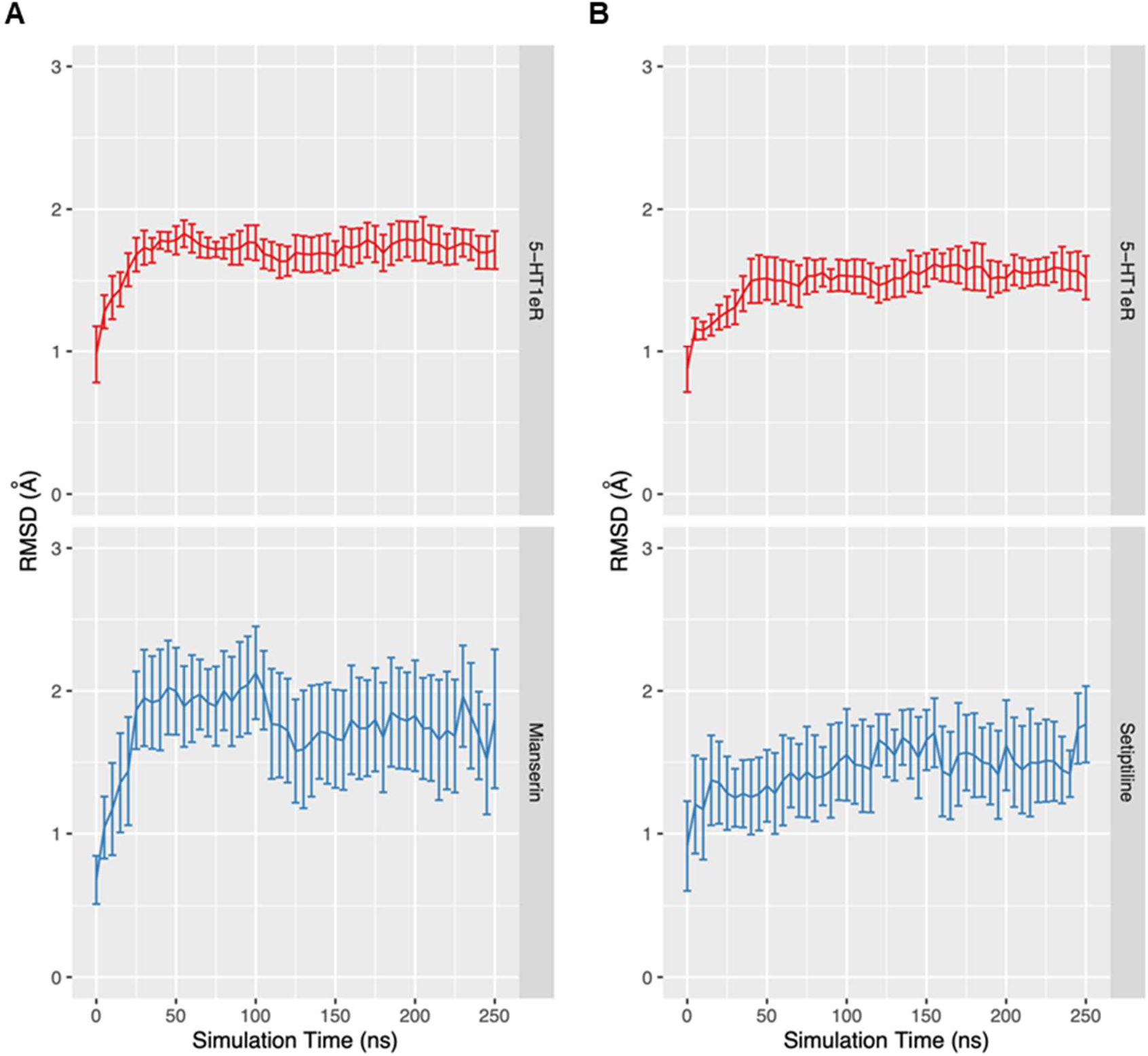
Molecular dynamics simulations of the mianserin- and setiptiline bound 5-HT_1e_R-G_i1_ complexes. Time evolution of the RMSD during four 250 ns production simulations of (**A**) the mianserin-bound 5-HT_1e_R heavy atoms (in red) and the mianserin heavy atoms (in blue) and (**B**) the setiptiline-bound 5-HT_1e_R heavy atoms (in red) and the setiptiline heavy atoms (in blue). All values are calculated after alignment of the 5-HT_1e_R heavy atoms. The average over the four 250 ns production simulations is reported with error bars referring to the 25th and 75th percentiles.

**Fig. S7.**
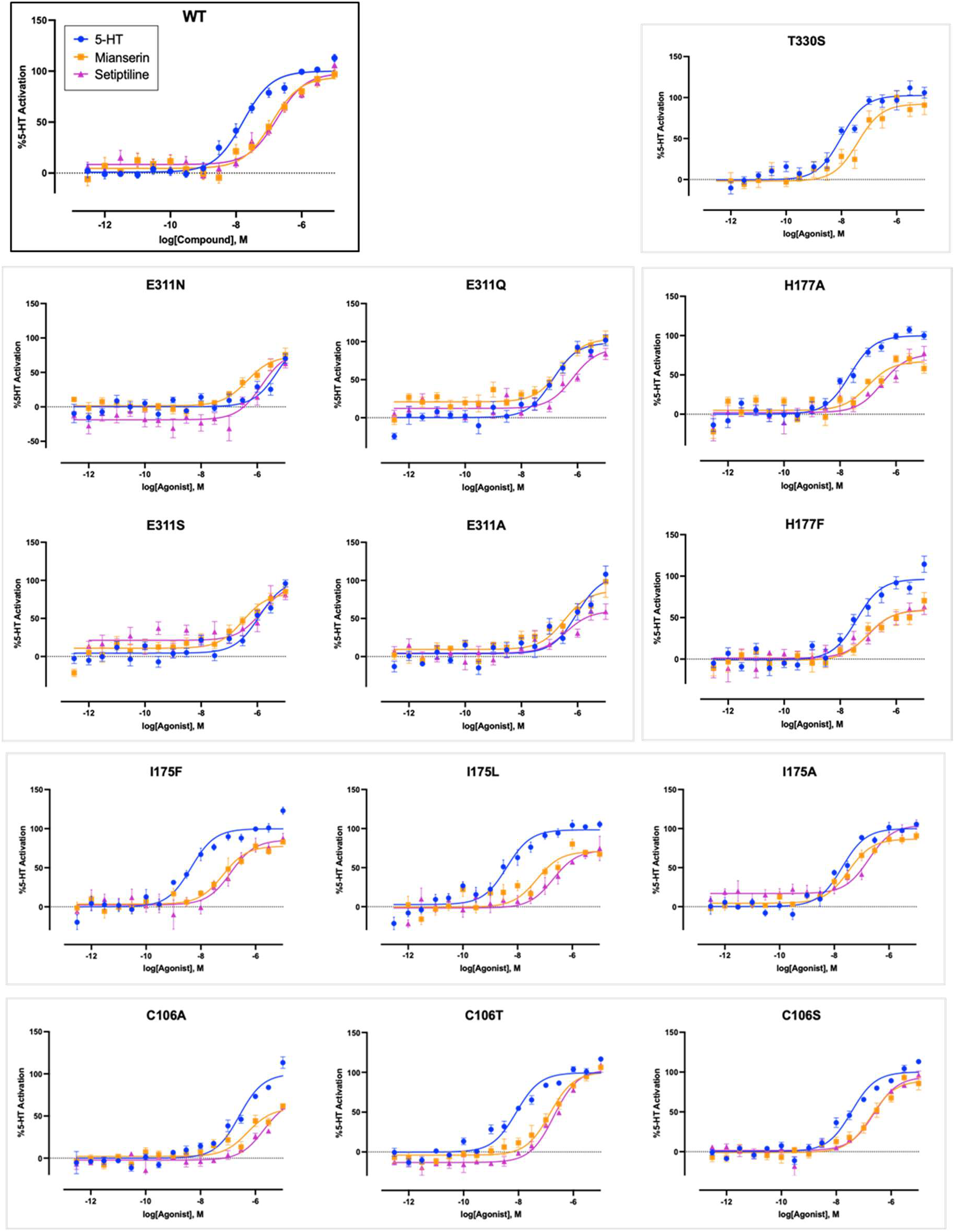
Effect of binding site mutations on the 5-HT_1e_R activities of 5-HT, mianserin, and setiptiline. Concentration response experiments determining the activity of 5-HT, mianserin, and setiptiline at wildtype (wt) and mutant 5-HT_1e_R as measured by G_i1_ BRET in HEK293T cells. Experiments were performed in triplicate and mean±SEM from at least two (n=2) independent experiments were averaged and normalized to 5-HT’s response.

**Fig. S8.**
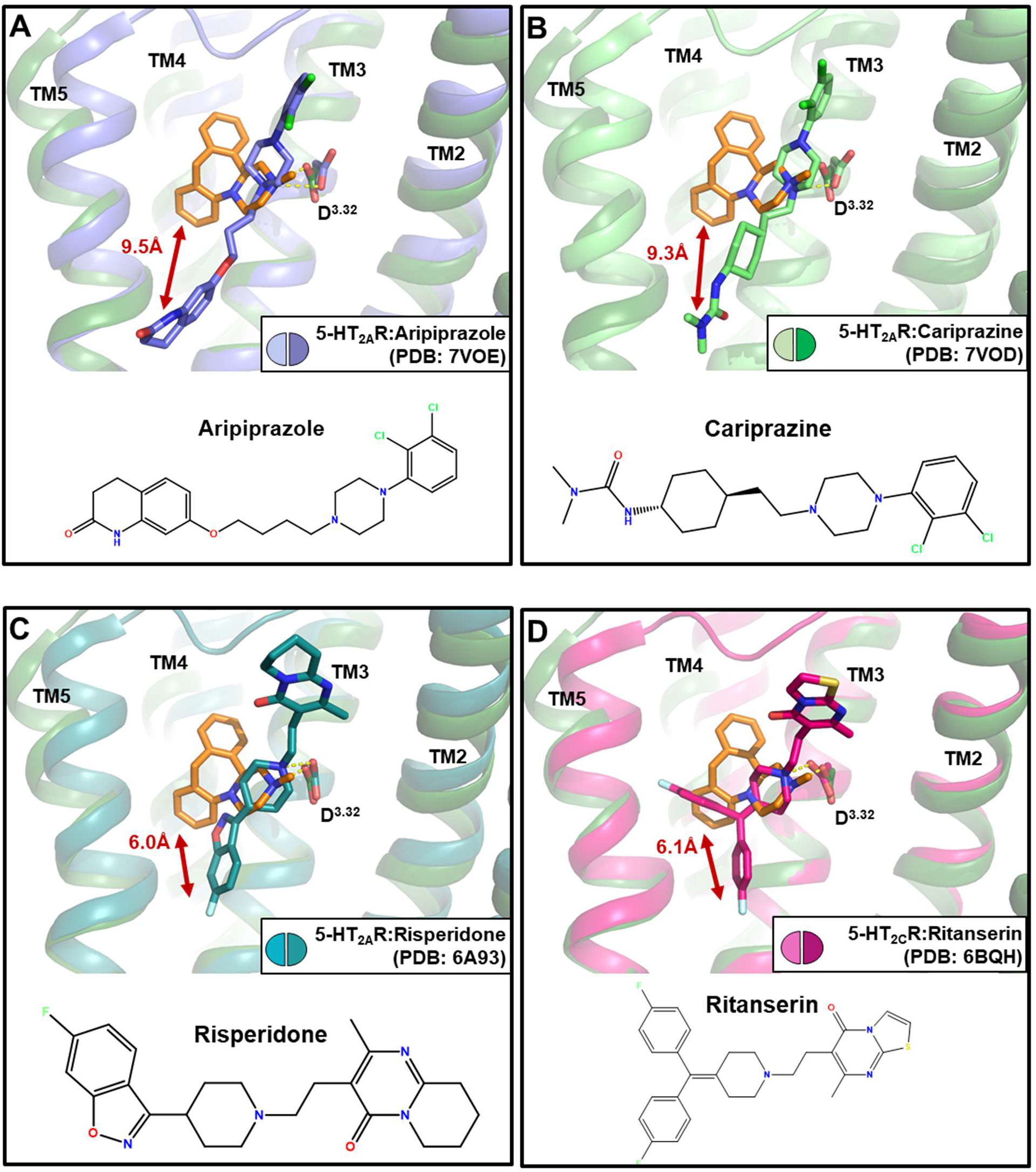
Structural comparison of mianserin-bound 5-HT_1e_R and inactive state 5-HT receptors bound to various antagonists. Superposition of mianserin-bound 5-HT_1e_R-G_i1_ complex structure and structures of aripiprazole-bound 5-HT_2A_R (**A**, blue, PDB: 7VOE), cariprazine-bound 5-HT_2A_R (**B**, green, PDB: 7VOD), risperidone-bound 5-HT_2A_R (**C**, teal, PDB: 6A93), and ritanserin-bound 5-HT_2C_R (**D**, pink, PDB: 6BQH). Ionic bonds between compounds and conserved D3.32 are indicated as dashed yellow lines. Distances between mianserin and antagonist drugs are measured between atoms closest to the receptor 7TM core and shown as red arrows.

**Table S1.**
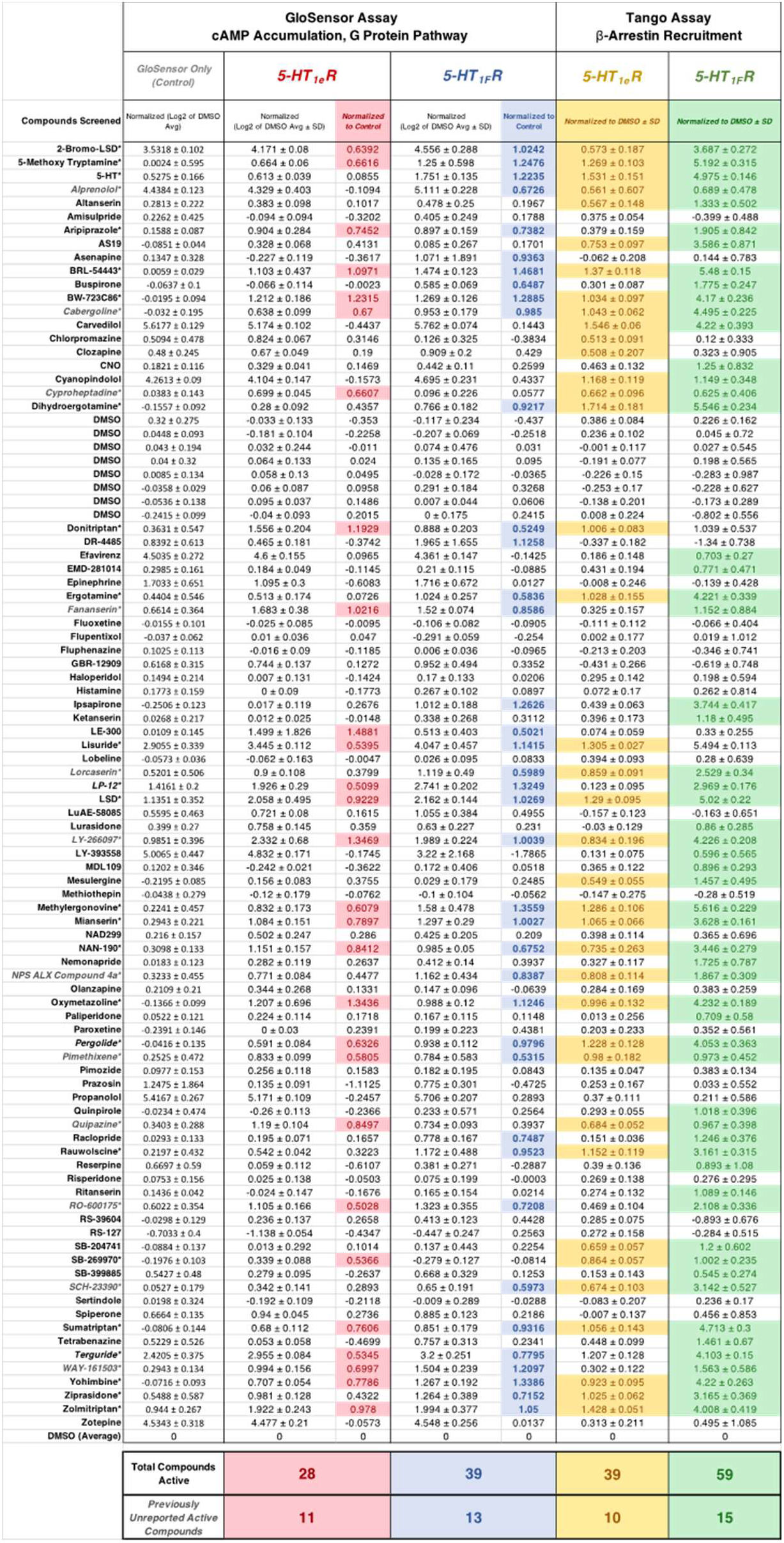
Activity of 10 µM aminergic drugs at 5-HT_1e_R and 5-HT_1F_R as determined by cAMP accumulation and arrestin-recruitment in HEK293T cells. Compounds that met the activation threshold of 0.5*log_2_FC over baseline are denoted with asterisks. Compounds not previously reported to display activity at 5-HT_1e_R and 5-HT_1F_R are denoted with italics and gray text.

**Table S2.**
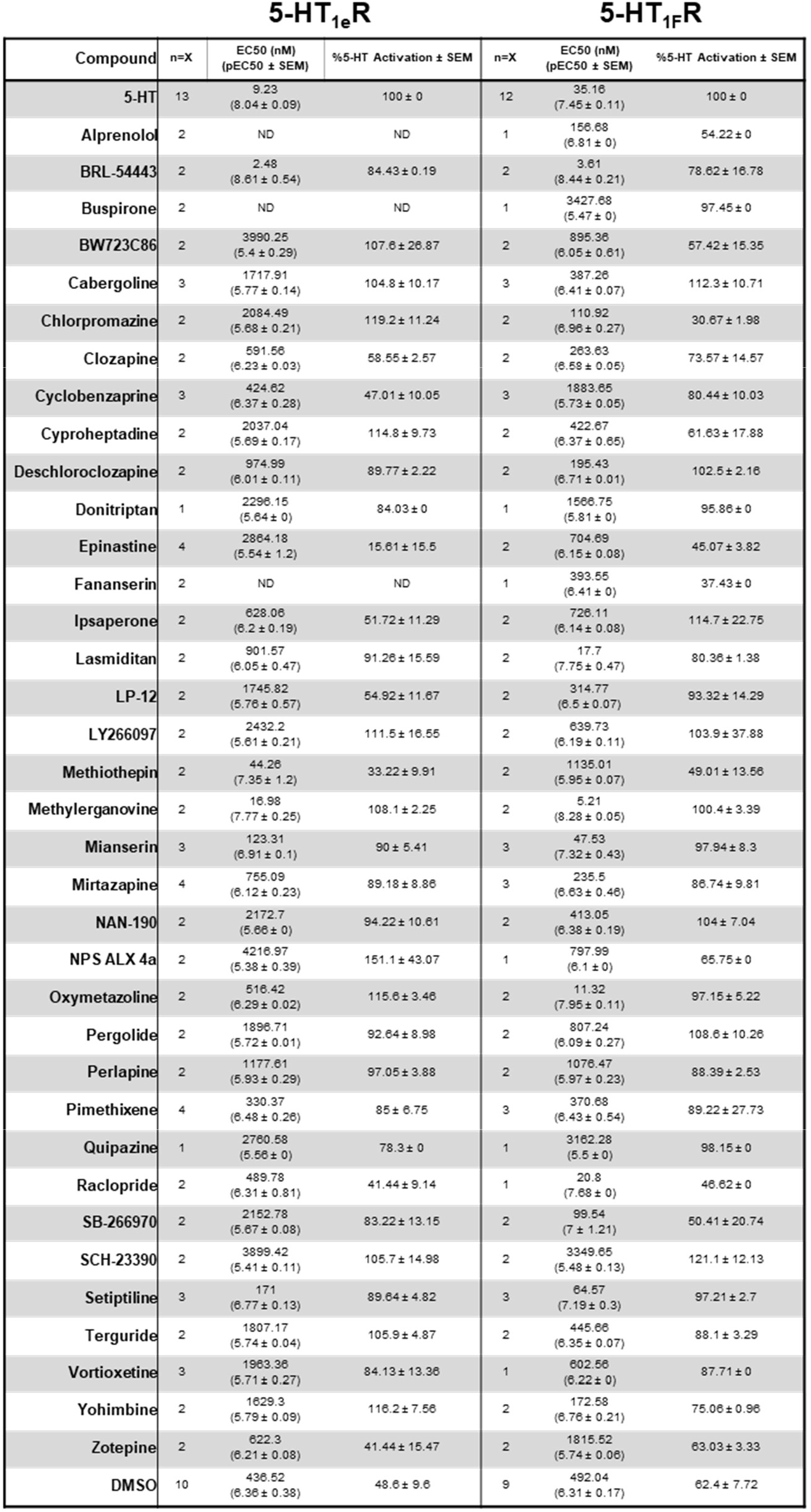
Potency and efficacy of aminergic drugs at 5-HT_1e_R and 5-HT_1F_R as determined by G_i1_ BRET G protein dissociation assays in HEK293T cells.

**Table S3.**
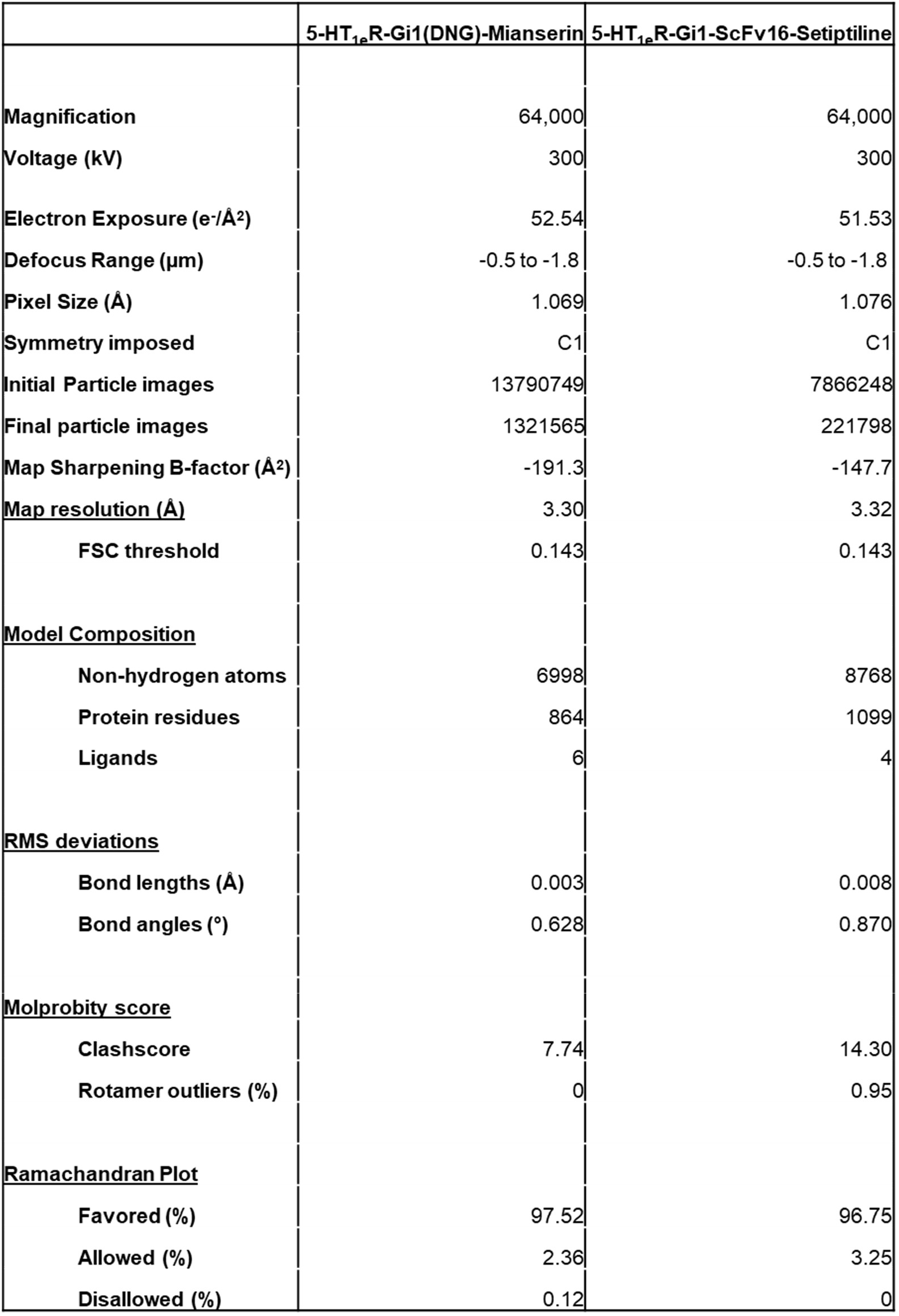
CryoEM data collection, refinement, and validation statistics.

**Table S4.** Activity of 5-HT, mianserin, and setiptiline at wildtype and mutant 5-HT_1e_R as determined by G_i1_ BRET G protein dissociation assays in HEK293T cells.

|  | 5-HT |  |  | Mianserin |  |  | Setiptiline |  |  |
| --- | --- | --- | --- | --- | --- | --- | --- | --- | --- |
|  | n=X | EC50 (nM)<br>(pEC50 ± SEM) | % 5-HT Activation ± SEM | n=X | EC50 (nM)<br>(pEC50 ± SEM) | % 5-HT Activation ± SEM | n=X | EC50 (nM)<br>(pEC50 ± SEM) | % 5-HT Activation ± SEM |
| E311A | 2 | 1936.42<br>(5.71 ± 0.03) | 100 ± 0 | 2 | 262.42<br>(6.58 ± 0.1) | 49.36 ± 1.34 | 2 | 130.02<br>(6.89 ± 0.94) | 59.49 ± 26.98 |
| E311S | 3 | 990.83<br>(6 ± 0.19) | 100 ± 0 | 3 | 220.29<br>(6.66 ± 0.19) | 74.09 ± 7.19 | 2 | 427.56<br>(6.37 ± 0.5) | 57.91 ± 0.19 |
| E311N | 2 | 4954.5<br>(5.31 ± 0.17) | 100 ± 0 | 2 | 440.55<br>(6.36 ± 0.06) | 71.88 ± 14.01 | 2 | 1172.2<br>(5.93 ± 0.15) | 98.94 ± 38.09 |
| E311Q | 3 | 133.35<br>(6.88 ± 0.22) | 100 ± 0 | 3 | 231.74<br>(6.64 ± 0.08) | 83.29 ± 4.33 | 3 | 534.56<br>(6.27 ± 0.33) | 81.25 ± 10.6 |
| E311D | 2 | 176.2<br>(6.75 ± 0.11) | 100 ± 0 | 2 | 89.74<br>(7.05 ± 0.26) | 90.43 ± 5.96 | 2 | 78.16<br>(7.11 ± 0.25) | 82.69 ± 4.93 |
| I175A | 3 | 18.58<br>(7.73 ± 0.09) | 100 ± 0 | 3 | 33.57<br>(7.47 ± 0.04) | 82.41 ± 1.12 | 2 | 186.64<br>(6.73 ± 0.04) | 87.81 ± 0.71 |
| I175F | 3 | 4.44<br>(8.35 ± 0.07) | 100 ± 0 | 3 | 65.61<br>(7.18 ± 0.04) | 74.54 ± 8.68 | 2 | 118.03<br>(6.93 ± 0.25) | 83.63 ± 0.4 |
| I175L | 4 | 3.49<br>(8.46 ± 0.39) | 100 ± 0 | 4 | 8.59<br>(8.07 ± 0.62) | 73.69 ± 10.47 | 2 | 199.53<br>(6.7 ± 0.05) | 78.48 ± 3.92 |
| I175W | 2 | 7.06<br>(8.15 ± 0.14) | 100 ± 0 | 2 | 7.48<br>(8.13 ± 0.06) | 88.55 ± 1.88 | 2 | 15<br>(7.82 ± 0.11) | 94.42 ± 2.86 |
| T330F | 2 | 0.09<br>(10.06 ± 2.19) | 100 ± 0 | 2 | 274.16<br>(6.56 ± 0.2) | 62.91 ± 17.14 | 0 | ND | ND |
| T330S | 2 | 10.64<br>(7.97 ± 0.21) | 100 ± 0 | 2 | 46.13<br>(7.34 ± 0.15) | 97.5 ± 5.56 | 0 | ND | ND |
| T330V | 2 | 82.79<br>(7.08 ± 0.01) | 100 ± 0 | 2 | 550.81<br>(6.26 ± 0.18) | 64.43 ± 1.96 | 2 | 579.43<br>(6.24 ± 0) | 62.09 ± 4.69 |
| H177A | 2 | 22.49<br>(7.65 ± 0.12) | 100 ± 0 | 2 | 99.54<br>(7 ± 0) | 62.24 ± 0.14 | 2 | 220.8<br>(6.66 ± 0.33) | 73.12 ± 8.76 |
| H177F | 3 | 53.46<br>(7.27 ± 0.24) | 100 ± 0 | 3 | 87.3<br>(7.06 ± 0.16) | 60.52 ± 7.2 | 3 | 102.09<br>(6.99 ± 0.27) | 58.26 ± 5.93 |
| H177T | 3 | 64.27<br>(7.19 ± 0.13) | 100 ± 0 | 3 | 316.23<br>(6.5 ± 0.27) | 78.34 ± 15.46 | 3 | 207.01<br>(6.68 ± 0.06) | 51.19 ± 11.26 |

